# Engineering Transcriptional Interference through RNA Polymerase Processivity Control

**DOI:** 10.1101/2020.09.23.310730

**Authors:** Nolan J. O’Connor, Antoni E. Bordoy, Anushree Chatterjee

## Abstract

Antisense transcription is widespread in all kingdoms of life and has been shown to influence gene expression through transcriptional interference (TI), a phenomenon in which one transcriptional process negatively influences another *in cis*. The processivity, or uninterrupted transcription, of an RNA Polymerase (RNAP) is closely tied to levels of antisense transcription in bacterial genomes, but its influence on TI, while likely important, is not well-characterized. Here we show that TI can be tuned through processivity control via three distinct antitermination strategies: the antibiotic bicyclomycin, phage protein Psu, and ribosome-RNAP coupling. We apply these methods toward TI and tune ribosome-RNAP coupling to produce 38-fold gene repression due to RNAP collisions. We then couple protein roadblock and RNAP collisions to design minimal genetic NAND and NOR logic gates. Together these results show the importance of processivity control for strong TI and demonstrate the potential for TI to create sophisticated switching responses.

## INTRODUCTION

Antisense transcription is widespread in all kingdoms of life. While once attributed largely to transcriptional noise from hidden or cryptic promoters (1), antisense transcription is now understood to govern import cellular decisions—for example, meiotic entry in *S. cerevisiae* (2), senescence effects in fibroblast cells (3), and antibiotic resistance plasmid conjugation in *E. faecalis* (4). More recently, high-resolution transcript mapping in bacteria has shown that antisense transcription delineates gene boundaries through bidirectional termination of transcription (5). Rho-dependent transcriptional termination is understood to suppress antisense transcription in bacteria (6), but antisense transcription has still been shown to regulate gene expression throughout the genome (5, 7).

There are two known modes of transcriptional regulation by antisense transcription: antisense RNA (asRNA) regulation, where sense and antisense RNAs hybridize to promote RNAse-mediated degradation or block the ribosome binding site to prevent its translation (8–10), and collisions of the transcriptional machinery originated from sense and antisense promoters, termed transcriptional interference (TI) (8, 10–12). Three primary modes of TI—RNA Polymerase (RNAP) collisions, sitting duck, and promoter occlusion—have been proposed (11) and parsed through experiments (13) and mathematical modeling (9, 14). Direct contact of bacterial RNAPs has not been observed during head-on RNAP collisions (15), and it is generally understood that interference of one RNAP on another may be mediated through DNA supercoiling (16, 17) rather than due to direct collisions of transcriptional machinery. Consequently, the act of cis-antisense transcription has been shown to reliably down-regulate gene expression (9, 13, 18–22).

While previous TI studies have thoroughly investigated the genetic architectures that influence the frequency of collisions—elements such as interfering and expressing promoter strength (18–20) and inter-promoter distance (20)—little attention has been paid to the number of transcription elongation factors that associate with RNAP during transcription. These proteins—such as NusG, which bridges an RNAP and ribosome during the pioneering round of transcription and, in the absence of a co-translating ribosome, facilitates Rho termination (6, 23, 24)—affect the processivity, i.e. the uninterrupted transcription, of an RNAP. Recent transcriptomic studies in bacteria have linked antisense transcription to Rho termination and the modulation of RNAP processivity (5–7, 25, 26), highlighting the importance of RNAP processivity to TI over protein coding sequences. For example, a head-on collision event between an ‘interfering’ RNAP transcribing an untranslated region and an ‘expressing’ RNAP that is coupled to a co-translating ribosome is likely biased toward the latter due to Rho termination of the former (Fig. 1a). We hypothesized that protecting interfering RNAPs from Rho termination through processivity control (Fig. 1a) could improve the strength of TI and enable its engineering for higher-order switching responses.

**Figure 1:**
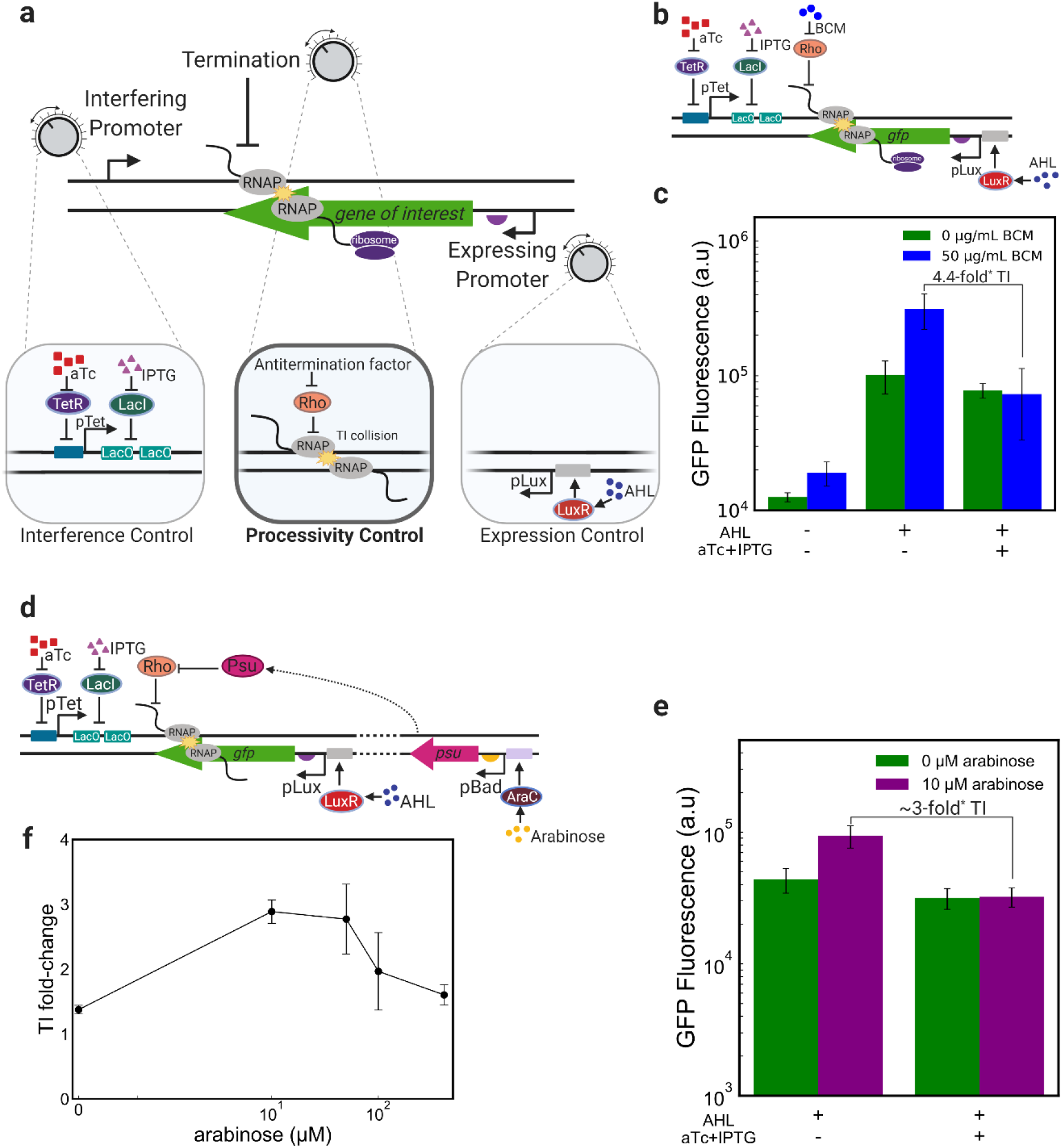
High processivity of interfering RNAPs is essential for strong TI. a) RNAP collision based transcriptional interference (TI) is tunable through the expression and interfering promoter strengths and RNAP processivity. Processivity control through antitermination of the interfering RNAP represents a novel strategy to engineer TI. b) Diagram showing the genetic elements comprising our inducible TI system and illustrating the effects of Rho and its inhibitor antibiotic, bicyclomycin (BCM) on the course of the interfering RNAP c) The addition of bicyclomycin (BCM) generates TI through the suppression of Rho termination of the interfering RNAP. d) Diagram of arabinose-inducible Rho inhibition via Psu. The protein Psu is under the control of the arabinose-inducible pBad promoter. e) Sublethal Psu expression with 10 μM of arabinose creates roughly 8-fold TI. In this experiment, AHL was present at a concentration of 100 μM, aTc was present at a concentration of 100 ng/mL and IPTG was present at a concentration of 1 mM. f) This construct shows a tunable TI system, with arabinose activating expression of Psu, which inhibits Rho and thereby improves the processivity of the interfering RNAP and strengthens TI. Error bars are denoted as ± s.d. Statistical significance as determined through the Mann-Whitney U test (p<0.05) denoted as *. n=3 biological replicates.

Here, we show that engineering processivity control of the interfering RNAP can tune TI. We demonstrate processivity control through the use of three antitermination mechanisms: the antibiotic bicyclomycin (27), expression of the phage polarity suppression protein Psu (28–30), and a co-translating ribosome (21, 31, 32) improve the strength of RNAP collisions (21, 23, 31). We engineer convergent gene constructs that permit an interfering RNAP-ribosome complex (‘expressome’(33)) to enter the opposing gene’s open reading frame, causing strong repression of gene expression, and creating, to our knowledge, the first synthetic expressome-on-expressome collision system. We show that processivity control, when coupled with control of interfering and expressing promoters (Fig. 1a) creates a layered, tunable TI system. We then apply these design rules to build two-input, minimal NAND and NOR transcriptional logic gates that couple protein roadblock with TI collisions. Together, our results demonstrate the importance of processivity control for tuning and engineering strong TI.

## MATERIALS AND METHODS

### Plasmids

Constructs containing fluorescent modules were cloned into pZE21MCS (Expressys) through restriction enzyme cloning and Gibson Assembly. SalI and BamHI were used for the insertion of GFP. GFP was obtained from pAKgfp1 (Addgene #14076). mCherry was obtained from PFPV-mCherry (Addgene #20956). BamHI and MluI were used to invert GFP for NAND and NOR constructs. ApaI was used to insert LuxR. NotI and AgeI were used to insert pLux. All restriction enzymes were purchased from Thermo Fisher. Insertion of *psu* was performed using a single-enzyme PciI digestion with FastAP. pBad promoters with araC were sourced from pX2_Cas9 and inserted using Gibson assembly. Primers for Gibson reactions are available upon request. The plasmid containing *Psu*, pHL 2067, was generously provided by Dr. Han Lim through Addgene.

Point mutations to introduce stop codons into antisense *mCherry* and create orthogonal araC* mutants were performed using a Quikchange (Agilent) PCR protocol. Single base-pair mismatches in forward and reverse primers were used in a modified PCR cycle to create mismatches, and DpnI was used to digest any original template.

### Strains and cell culture

Cloning and experiments to show logic behavior using TI with GFP and mCherry were performed in *E. coli* strain DH5αZ1 (Expressys). Transformation colonies were grown in Luria-Bertani (LB) and agar plates supplemented with kanamycin (50 μg/mL).

### GFP and mCherry induction assays

Individual colonies were picked from LB and agar plates supplemented 50 μg/mL kanamycin and incubated for 16 h at 37 °C under orbital shaking at 200 rpm. Then, the cells were diluted 1:10 into fresh LB media supplemented with 50 μg/mL kanamycin. Induction was performed at various inducer concentrations using anhydrous tetracycline (aTc), isopropyl β-D-1-thiogalactopyranoside (IPTG), 3-oxo-dodecanoyl-L-homoserine lactone (AHL). AHL powder was dissolved in a solution of 99.99% ethyl acetate and 0.01% glacial acetic acid, aliquoted as needed, and stored long term at −20 °C. Cells were grown for 6-8 h at 37 °C under shaking in a flat bottom 96-well plate in a microplate reader (Tecan Genios). Optical density at 590 nm was measured during induction. Following the growth period, the cells were transferred to a V-bottom 96-well plate and pelleted by centrifugation of the plate at 4000 rpm for 5 minutes at 4 °C. The supernatant was removed by vigorously inverting the plate and then the pellets were resuspended in 100 μL PBS+4% formaldehyde and transferred to a flat bottom plate, which was then stored at 4 °C prior to flow cytometry measurements.

#### Bicyclomycin (BCM) Treatments

50 ng/mL bicyclomycin (Santa Cruz Biotechnology) was added along with other inducers (aTc, IPTG, AHL) after 3 hours of growth under orbital shaking at 200 rpm in LB+Kan following a 1:10 dilution of overnight cultures grown for 16 hours. Cells were grown for an additional 3 hours before being washed and fixed in PBS+4% formaldehyde and subsequently measured with flow cytometry.

#### Psu Experiments

To mitigate the adverse growth effects of high Psu expression, inducers were added to microplate wells (to achieve a total volume of 100 μL) after 2 hours of growth under orbital shaking at 200 rpm in LB+Kan following a 1:10 dilution of overnight cultures grown for 16 hours. Cells were grown for an additional 4 hours before being washed and fixed in PBS+4% formaldehyde and subsequently measured with flow cytometry.

### Flow cytometry

Before fluorescence measurements conducted with a FACSCelesta instrument, samples were diluted 1:50 in PBS. The 588B 530/30V (800 V) channel was used to measure GFP levels. FSC-V=420 V, SSC-V=260 V, FSC-Threshold= 8000, SSC-Threshold= 200. For each sample, 50,000 cells were measured. At least four biological replicates were collected for each construct. Data was analyzed using FlowCytometryTools package in Python 3.7. Statistical differences were examined using the Mann-Whitney *U* test.

To calculate the TI fold-change for a particular construct or set of conditions, mean fluorescence values of biological replicates (minimum 3) were averaged and used in the following equation:

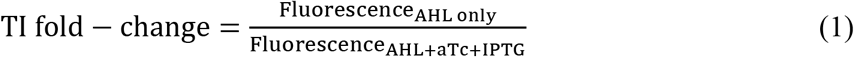

### Strand-specific qPCR

#### Growth experiment and RNA isolation

1 mL overnights in LB media supplemented with 50 μg/mL kanamycin were grown for at 37 °C under orbital shaking 16 hours. Overnight cultures were diluted 1:50 into LB media with 50 μg/mL kanamycin and relevant inducers (aTc, IPTG, AHL) and grown for 6 hours at 37 °C with orbital shaking at a total volume of 1.5 mL. Cell pellets were spun down for 2 minutes at 12000 rcf, supernatant was removed, and pellets were stored at −80 °C prior to RNA extraction. RNA was extracted and purified using GeneJET RNA Purification Kit (Thermo Fisher). Total RNA was digested with DNase I at 37 °C for 30 minutes and subsequently repurified.

#### cDNA synthesis

cDNA synthesis was carried out using High Capacity cDNA Reverse Transcription Kit (Applied Biosystems). Gene-specific primers corresponding to the antisense *mCherry* sequence were used to prime cDNA synthesis. 2 μL of primer at 10 μM and ~500 ng of RNA were added to the 24 μL reaction. cDNA synthesis was carried out using the temperature steps: 25 °C for 10 minutes; 37 °C for 2 hours, 85 C for 5 minutes, 4 °C hold.

#### RT-qPCR

RT-qPCR was carried out using FastStart Universal SYBR Green Master Mix (Rox) (Sigma Aldrich) in an Applied Biosystems QuantStudio 6. 1.5 μL of cDNA, 1 μL reverse and forward primers were added to each 10 μL reaction. *Kanamycin* was used as a reference housekeeping gene for each construct. Threshold values were normalized to *Kan* (ΔC_T_), and these ΔC_T_ values for *gfp-mCherry* and *gfp*-mCherry* (Fig. 2d) or no aTc+IPTG and aTc+IPTG (Fig. 2e) were compared (ΔΔC_T_). Error from biological replicates was propagated through the normalization and comparisons. Fold-change error bounds are reported as 2^−(ΔΔCT + sd)^ and 2^−(ΔΔCT - sd)^ calculated comparing the CT values for *gfp-mCherry* and *gfp*-mCherry* (Fig. 2d) or no aTc+IPTG to 100 ng/mL aTc + 1 mM IPTG (Fig. 2e).

**Figure 2:**
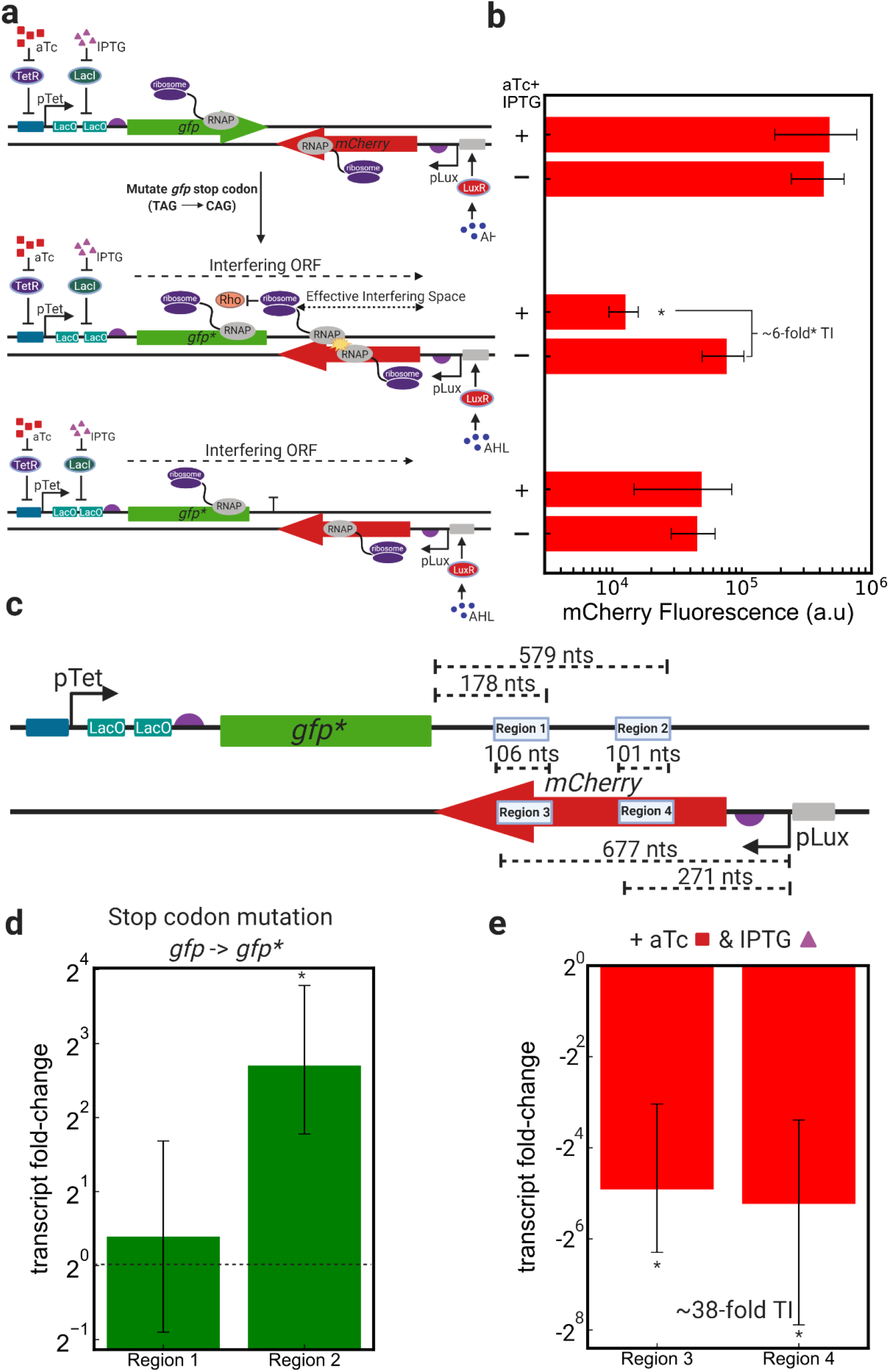
Protecting the interfering RNAP in an expressome complex improves processivity and creates strong TI. a) Designs of convergent *gfp-mCherry* constructs: (from top to bottom), convergent *gfp* and *mCherry* genes under the control of pTet+LacO and AHL, respectively; convergent *gfp* and *mCherry* genes with the *gfp* stop codon mutated to a glutamine (Gln) amino acid (denoted as *gfp\**), thereby extending the ORF originating from downstream of the pTet-LacO 5’-UTR through the mCherry ORF until 2 bp before the pLux transcription start site; a strong unidirectional terminator was inserted downstream of *gfp\** to impede the progress of the interfering expressome. b) mCherry expression with 120 μM AHL and with and without 100 ng/mL aTc and 1 mM IPTG demonstrates the effects of TI on mCherry expression in all constructs. c) Strand-specific qPCR was used to target transcripts containing regions of the antisense (Regions 1 and 2) and sense (Regions 3 and 4) *mCherry* transcripts. Regions 1 and 3 and Regions 2 and 4 represent the same amplicons, of sizes 106 and 101 nts, respectively, with different gene-specific cDNA priming (Materials and Methods). d) Measuring antisense *mCherry* transcript fold-change (using *Kan* as a reference) upon *gfp* stop codon mutation (comparing *gfp-mCherry to gfp*-mCherry*) shows improved processivity (increase in Region 2 transcripts) upon ribosome-RNAP coupling. Cells containing both constructs were grown in the absence of any AHL (no pLux activation) and in the presence of 100 ng/mL aTc and 1 mM IPTG, in order to measure processivity of the interfering RNAP. Data titled ‘Region 1’ and ‘Region 2’ represent transcripts that contain those amplicon regions (Materials and Methods). d) Measuring sense *mCherry* transcript fold-change (using *Kan* as a reference) upon interfering promoter activation (comparing AHL-only condition to AHL with aTc+IPTG) shows ~38-fold TI (decrease in Regions 3 and 4 transcripts) upon interfering promoter activation. Cells containing both constructs were grown with 200 μM AHL (full pLux activation) and in the presence or absence of 100 ng/mL aTc and 1 mM IPTG in order to measure knockdown of the *mCherry* transcript due to TI. Fold-change represents reduction in transcript levels upon the activation of the interfering promoter, pTet, with aTc and IPTG. Data titled ‘Region 3’ and ‘Region 4’ represent transcripts that contain those amplicon regions (Materials and Methods). For figures d and e: fold-change represents changes in transcript levels (2^−ΔΔCT^) upon either mutation of stop codon (d) or induction of interfering promoter (e). Error bars are denoted as ± s.d, in (d-e) represented as (2^−(ΔΔCT+s.d)^, 2^−(ΔΔCT-s.d)^). * indicates significance (Mann-Whitney U-test (b), one-sample T-test (d-e), p<0.05) in the expression differences of induced vs. uninduced interfering promoter (b) or of the ΔΔC_T_ values with respect the null-hypothesis of ΔΔC_T_=0 d-e). n=3 biological replicates.

## RESULTS

### Processivity control is essential for strong TI

Transcriptional Interference (TI) resulting from convergent promoters has been shown to depend on levels of interference and expression control through deliberate tuning of promoter strengths (9, 18–20) (Fig. 1a). Here we chose to use an inducible promoter system in order to more easily tune their relative strengths. The quorum sensing promoter pLux, induced with AHL, is used as the “expression control” module to regulate *gfp* production. The aTc-inducible pTet is oriented antisense to *gfp* and is used as the “interference control” module to downregulate *gfp* expression through antisense transcription. We added an IPTG-inducible lac operator 47 bp downstream of the pTet transcription start site (Fig. 1a) to further tune the interference control via protein roadblock (34). (DNA lengths for the relevant plasmids are illustrated in Supplementary Figure S1.)

We observed that at highest strength of interference, achieved when this pTet-LacO architecture (Fig. 1b) was activated with saturating aTc and IPTG in order to fire interfering RNAPs to repress GFP expression from pLux, a significant but weak ~1.6-fold change in GFP expression occurred (Equation 1, Supplementary Figure S2). The extent of this TI repression was dependent on both pTet and pLux activity (aTc and AHL concentration, respectively), decreasing with high AHL concentrations and increasing with high aTc concentrations (Supplementary Figure S2). These results demonstrate the effects of interference and expression control on TI and generally agrees with other TI studies, which have found that a strong interfering promoter and weak expressing promoter are required to produce significant TI (18–20).

We posited that the lack of observed strong repression from TI may be related to low processivity of the interfering RNAP. *In vivo*, elongating RNAPs interact with transcriptional factors that modulate its processivity (23). For example, the protein NusG associates with elongating RNAPs to prevent pausing and backtracking (6, 31) and aids Rho in factor-dependent termination (6) of RNAPs transcribing untranslated mRNA. Rho primarily targets RNAPs transcribing in untranslated regions of DNA in order to suppress pervasive transcription in the genome (6, 21). Because the mRNA transcribed by the interfering RNAP in this construct is not simultaneously being translated, it is susceptible to Rho termination, which may explain the relatively low levels of observed interference.

To test this hypothesis, we exposed exponentially growing cells to a sub-lethal dose of bicyclomycin (BCM), an antibiotic that targets the ATP turnover of Rho, thereby alleviating factor-dependent termination(35) (Fig. 1a). The interaction of Rho with BCM provides the “processivity control” in the construct. Upon BCM addition we observed a significant 4.4-fold increase in TI compared to no treatment (Fig. 1c), suggesting that Rho inhibition increases interfering RNAP processivity and allows for strong TI. Note that the condition-wide increase in GFP expression in the presence of BCM likely results from decreased termination in the 32 bp 5’ UTR region of the expressing promoter. Importantly, the difference in expression between the AHL-only and AHL+aTc+IPTG (Fig. 1c) indicates an improvement in interference for RNAPs originating from pTet. These results suggest that the extent of TI can be tuned through control of RNAP processivity.

### Phage polarity suppression protein Psu tunes TI

The manipulation of RNAP processivity using BCM suggested that the strength of RNAP collisions can be controlled through inhibition of Rho activity. To further fine-tune Rho inibition, we incorporated the P4 phage protein *Psu* into our plasmid, under a pBad promoter (Supplementary Figure S1, Fig. 1d). Similar to BCM, Psu prevents Rho from translocating along the nascent mRNA through inhibition of ATP hydrolysis (28) and has previously been shown to improve RNAP processivity in *E. coli* (30). To our knowledge, Psu has never before been used to study TI. To reduce crosstalk between IPTG-inducible LacO and arabinose-inducible pBad, we used an araC mutant evolved to respond only to arabinose (36).

We found that, like BCM, arabinose-induced Psu expression both increases GFP expression and increases the fold-change in TI here to nearly 3-fold (Fig. 1e). We also found that induction of *Psu* with arabinose changed the extent of TI in a dose-dependent manner (Fig. 1f), representing a tunable TI system. Interestingly, high levels of Psu induction decreased the observed levels of TI, possibly due to large overall increases in protein expression and promoter leakiness, or a global disruption in gene expression (Supplementary Figure S3). We found that toxicity of Psu expression was neglible if Psu expression was induced after 2 hours of growth under orbital shaking (OD of ~0.3) (Supplementary Figure S4). Growth effects were, however, observed when arabinose was added upon dilution from overnights, at t=0 (Materials and Methods).

Both BCM and Psu have previously been shown to increase transcript production for genes that were susceptible to Rho-dependent termination while leaving protein levels unchanged, likely due to exclusion of the ribosome (30) resulting from BCM or Psu locking Rho in place on the transcript (29). The observed increase in fluorescence in our system could be dependent on the 5’ UTR of the fluorescent reporter, as a change in length and sequence in this region may reduce interference between Rho and the ribosome.

### Ribosomal protection of the interfering RNAP enhances TI over a gene of interest

The use of BCM and Psu disrupt Rho termination throughout the cell, limiting their applicability as processivity control strategies. We therefore sought a way to control RNAP processivity in only a gene of interest. The ribosome, when coupled with a transcribing RNAP, protects that RNAP from Rho termination, either through blockage of rho utilization sites and/or by sequestering NusG through interactions with the NusG CTD and S10 ribosomal subunit (6). Recently, direct interactions of a bacterial RNAP and ribosome have also been reported and termed the ‘expressome’ (33). Though there exists evidence for both bridged and direct contact between the RNAP and the ribosome and it is unclear when one linkage might occur (31), we will heretofore refer to the RNAP-ribosome complex as the ‘expressome’. It has been shown that co-translation of ribosomes along with elongating RNAPs can prevent the pre-mature termination of the latter by precluding Rho binding (21, 23, 31, 32). Protecting the interfering RNAP from Rho termination with a co-translating ribosome in an expressome complex should therefore strengthen gene repression through TI. To this end, we created a construct of convergently oriented *gfp* and *mCherry* sequences under the control of pTet-LacO and pLux, respectively (Fig. 2a, top). At high AHL concentrations, the release of interfering RNAPs using saturating aTc and IPTG did not significantly change mCherry expression (Fig. 2b, top). Interestingly, we did observe substantial TI of GFP as a function of the interfering and expressing promoter strengths (Supplementary Figure S5), which may result from sequence differences between the two fluorescent proteins, either in the form of pause sites or Rho utilization sites.

In this convergent *gfp-mCherry* construct, the interfering RNAP has already decoupled from its co-translating ribosome before it transcribes into the *mCherry* ORF (Fig. 2a). This ‘naked’ interfering RNAP is more exposed to Rho termination than the expressome complex, and this de-coupling of RNAP and ribosome likely explains the lack of observed TI at high AHL concentrations (Fig. 2b, top). We hypothesized that if we could prevent RNAP-ribosome de-coupling and allow the expressome enter to the *mCherry* ORF, the resulting increase in interfering RNAP processivity might strengthen TI repression of mCherry. To test this hypothesis, we mutated the stop codon of *gfp* to extend the open reading frame (ORF) of the interfering expressome into the *mCherry* ORF (Fig. 2a, middle). (Note that the notation change of *gfp to gfp\** reflects a complete abolition of GFP expression.) This point mutation resulted in significant, ~6-fold gene repression (Fig. 2b, middle) due to the improved processivity of the interfering RNAP when coupled with a co-translating ribosome. Interestingly, the mutation of the *gfp* stop codon created an ‘interfering ORF’ that extended through the antisense *mCherry* ORF and did not encounter an in-frame stop codon until 2 bp prior to the expressing promoter, pLux. Such a long ‘effective interfering space’ (Fig. 2a, middle) likely contributed to the improved strength of ribosome-coupled interfering RNAPs.

To confirm that the observed reduction in *mCherry* upon activation of pTet-LacO was due to TI, we added a strong unidirectional terminator (rrnBT1(37)) on the *gfp\** strand between the *gfp\** and *mCherry* sequences (Fig. 2a, bottom) in order to block interfering expressomes from entering the mCherry ORF. Note that this terminator does not introduce any stop codons into the interfering ORF and therefore maintains the interfering expressome course required for strong repression in this construct (Fig. 2b). We observed no significant TI when the interfering pTet-LacO module was induced with saturating aTc and IPTG (Fig. 2b, bottom), indicating that interactions between transcriptional machinery are likely responsible for the observed gene repression in the *gfp*-mCherry* construct. These results suggest that ribosome-aided RNAP processivity can create strong TI over a gene of interest.

To confirm that the *gfp* stop codon mutation improved RNAP processivity, we used strand-specific quantitative PCR (qPCR, Materials and Methods) to measure the abundance of transcripts antisense to *mCherry* in constructs with and without a *gfp* stop codon (Fig. 2c). Under saturating aTc and IPTG and with no AHL, we measured the relative amounts of transcripts that were long enough to contain regions 1 and 2, located 178 and 579 nts from the 3’ end of *gfp* or *gfp\**, respectively. These data showed that mutating the *gfp* stop codon does not significantly change the abundance of transcripts long enough to contain region 1 but does significantly change the abundance of transcripts containing region 2, at a 6.5-fold increase (Fig. 2d). This increase in long antisense transcripts provides transcription-level evidence that the *gfp* stop codon mutation (Fig. 2a) improves processivity of the interfering RNAP, as the interfering RNAP, when coupled to a ribosome, is able to transcribe further into the *mCherry* ORF on the antisense strand. This suggests that the TI observed measuring mCherry protein levels (Fig. 2b) can be attributed to improved interfering RNAP processivity.

The TI resulting from this improved processivity of the interfering RNAP is also evident when measuring levels of *mCherry* transcript. Measuring the relative abundance of the *mCherry* transcript with saturating AHL and in the presence and absence of interfering promoter induction (with saturating aTc and IPTG) demonstrates significant, ~38-fold knockdown of the *mCherry* transcript due to TI (Fig. 2e). Amplicons on the 5’ and 3’ ends of the *mCherry* transcript, regions 3 and 4 (Fig. 2c) were uniformly downregulated upon induction of the interfering promoter. Interestingly, in both the presence and absence of interfering promoter induction, there are ~13-fold more transcripts containing only region 3 than there are transcripts containing regions 3 and 4 (Supplementary Figure S6). This suggests that a number of truncated *mCherry* transcripts are produced in the *gfp*-mCherry* independent of TI. Both regions are downregulated upon interfering promoter induction, suggesting TI-induced knockdown, but the ratio between the abundances of each transcript length is maintained (Supplementary Figure S6). Premature transcriptional termination has been reported and is a function of the 5’ UTR sequence and secondary structure and RBS strength (30). It is surprising, though, that TI does not affect the relative amounts of truncated transcripts, given that TI is known to create truncated transcripts (4). This result suggests that TI collisions occur upstream of region 3, toward pLux, or that TI produces truncated *mCherry* transcripts that maintain the ~13-fold difference between short and long mRNA.

### TI from ribosome-RNAP coupling is tunable through promoter control

Previous TI studies have shown that the strength of RNAP collisions is a function of promoter strength (19–21). Likewise, here we find that TI in this ribosome-aided system can also be tuned through the activation of both expressing (pLux) and interfering (pTet) promoters (Figure Supplementary Figure S7), demonstrating a layered response to processivity, expression, and interference control (Fig. 1a). Previous TI studies have demonstrated an inverse relationship between activating promoter strength and TI (19–21), but interestingly here we observed TI fold-change—the ratio of fluorescence observed when the interfering promoter is induced vs. uninduced (See Materials and Methods, Equation 1)—unchanged for AHL concentrations greater than 20 μM (Supplementary Figure S8). In the absence of a LacI roadblock (at 1mM IPTG), increasing aTc concentrations reduce mCherry expression due to RNAP collisions, even at high AHL concentrations (Supplementary Figure S7). The aTc-dependence and the dependence of ribosome co-translation on TI fold-change demonstrates the importance of collision location: RNAP collisions must occur over the *mCherry* ORF—an ‘effective interfering space’ (Fig. 2a, middle)—in order to result in observable interference in this system. The coupled effects of strong interfering promoter strength (high aTc) and interfering RNAP processivity (ribosome-RNAP coupling) enable strong TI.

### TI strength is a function of the interfering expressome’s ORF length

The strong TI observed when the interfering expressome is allowed to read far into the *mCherry* ORF (Fig. 2a-b, middle; Fig 2d), when contrasted with the absence of TI when the interfering expressome encountered a stop codon at the end of *gfp* (Fig. 2a-b, top; Fig 2d) suggests the importance to the length of the interfering ORF. Because the likelihood of termination increases after the interfering RNAP decouples from the translating ribosome (21), the effect of TI should be stronger the longer the RNAP and ribosome can stay coupled.

We tested this hypothesis by introducing stop codons into the antisense *mCherry* sequence, using codon degeneracy to retain the mCherry amino acid sequence (Fig. 3a, Supplementary Table S3). These stop codons effectively changed the length of effective interfering space from 744 nts (distance from end of *gfp\** to stop codon, for the construct shown in Fig. 2) down to lengths of 36, 201, 369, and 639 nts, and up to 834 nts. Transcription and translation therefore uncoupled at different points along the mCherry ORF (Fig. 3a). The positive correlation between interfering ORF length and TI fold-change (Pearson correlation coefficient=0.97, p-value<0.05) shows that the location of this uncoupling influences the extent of TI (Fig. 3b, right), with early stop codons introduced into the *mCherry* antisense sequence nearly abolishing TI and stop codons downstream of pLux achieving ~10-fold TI. This result also demonstrates that processivity control through RNAP-ribosome coupling is tunable. Notably, there was barely a trend in TI when the interfering ORF extended past the *mCherry* sequence, at approximately 700 nts, suggesting that pLux promoter occlusion was minimal here. This result agrees with previous mathematical modeling studies (9, 13), showing that RNAP collision is the dominant form of TI over large intergenic regions.

**Figure 3:**
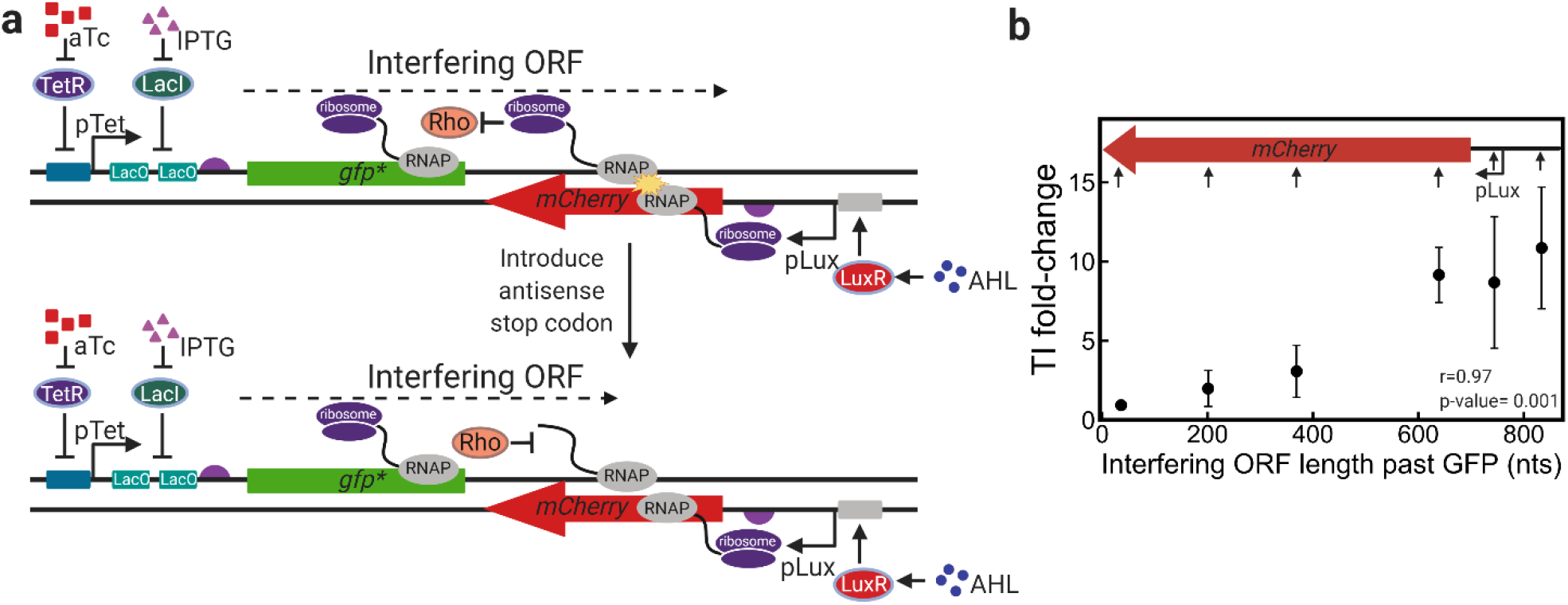
TI from a ribosome-protected interfering RNAP is tunable. Using codon degeneracy, stop codons were introduce in the mCherry antisense sequence, maintaining the mCherry amino acid sequence. a) An example illustrating how these stop codon-introducting point mutations shorten the ‘interfering ORF’ of the interfering expressome. When the interfering ORF is shorter, Rho has a higher chance of terminating transcription of the interfering RNAP and reducing the amount of observed TI. b) Measuring TI for each construct at identical AHL, aTc, and IPTG concentrations, a trend emerges in which the length of the ORF starting from the start codon of *gfp\** within the *mCherry* gene dictates the extent of TI. The significant (p-value<0.05) Pearson correlation coefficient suggests a positive relationship between interering ORF length and TI fold-change. The inserted pLux - *mCherry* region at the top of the figure shows the positions of the stop codons (represented here as upward arrows) introduced into the antisense strand. The x-axis denotes how many nts the interfering expressome will read before encountering a stop codon. Error bars are denoted as ± s.d with n≥4 biological replicates.

Reductions in gene expression due to convergently-oriented promoters are composed of both collisions of transcriptional machinery and interactions of sense and antisense RNA(8, 10, 11). Previous studies have knocked out promoters to prevent collisions of transcriptional machinery on the same strand and have found, in some cases significant asRNA intereference (19, 21) or neglible asRNA interference (18). Here, we have demonstrated that gene repression due to TI can be reliably tuned through modulation of interfering RNAP processivity, through the introduction of stop codons or terminators that prevent the interfering expressome from reading into the mCherry ORF. These results suggest that collisions, not asRNA interference, are the dominant mechanism, as asRNA likely would not experience such an interfering ORF length dependence. (Further discussion on the contributions in this system of TI and asRNA interference in this system are discussed further in Supplemental Note 1.)

### Ribosome-protected TI and roadblock together can produce NAND/NOR logic behaviors

We next sought to apply processivity control to engineer TI. A handful of prior studies have applied TI to create one-input logic gates (19), tuning of genetic switches (18, 19), and positive selection systems (20), and most recently control of metabolism genes in the *E. Coli* genome (38). Given that gene regulation via TI uses a low genetic footprint, TI-based genetic devices may be advantageous to the design of larger, more complex genetic programs (34). Further, coupling processivity control with interference and expression control (Fig. 1a) produces a layered response with several ‘knobs’ to tune that adjust the strength or behavior of a TI-based circuit. We therefore proposed that TI with processivity control could be used to design higher-order genetic circuits, such as two-input logic gates.

We recently demonstrated that TI from an inducible promoter upstream of an inducible roadblock can be rationally engineered to produce AND logic behavior responsive to two chemical inducers, aTc and IPTG (34). We also demonstrated that replacing this roadblock with an inducible promoter creates OR logic behavior after increasing the KD of the LacI roadblock in order to allow some readthrough from the upstream promoter while lowering leaky expression from pLac. It follows then that the logic modules used to express AND and OR-like behaviors can be used to control the release of RNAPs that represses GFP expression through RNAP collisions, effectively inverting the logic from AND/OR to NOT AND/OR, i.e. NAND/NOR.

To create NAND behavior, we used the inducible promoter pTet and dowstream protein roadblock LacI to control the release of interfering RNAPs that suppress mCherry expression through collisions (Fig. 4a, left). The release of the RNAPs is governed by a two-input AND logic gate, with aTc and IPTG as the inputs, and the interfering expressomes reduce gene expression through collisions, thereby effectively layering a NOT gate onto an AND gate, yielding NAND logic behavior. Using the *gfp*-mCherry* system for ribosome-protected processivity control, we demonstrate good NAND behavior with a 7.8-fold reduction in gene expression due to collision (Fig. 4a, right).

**Figure 4:**
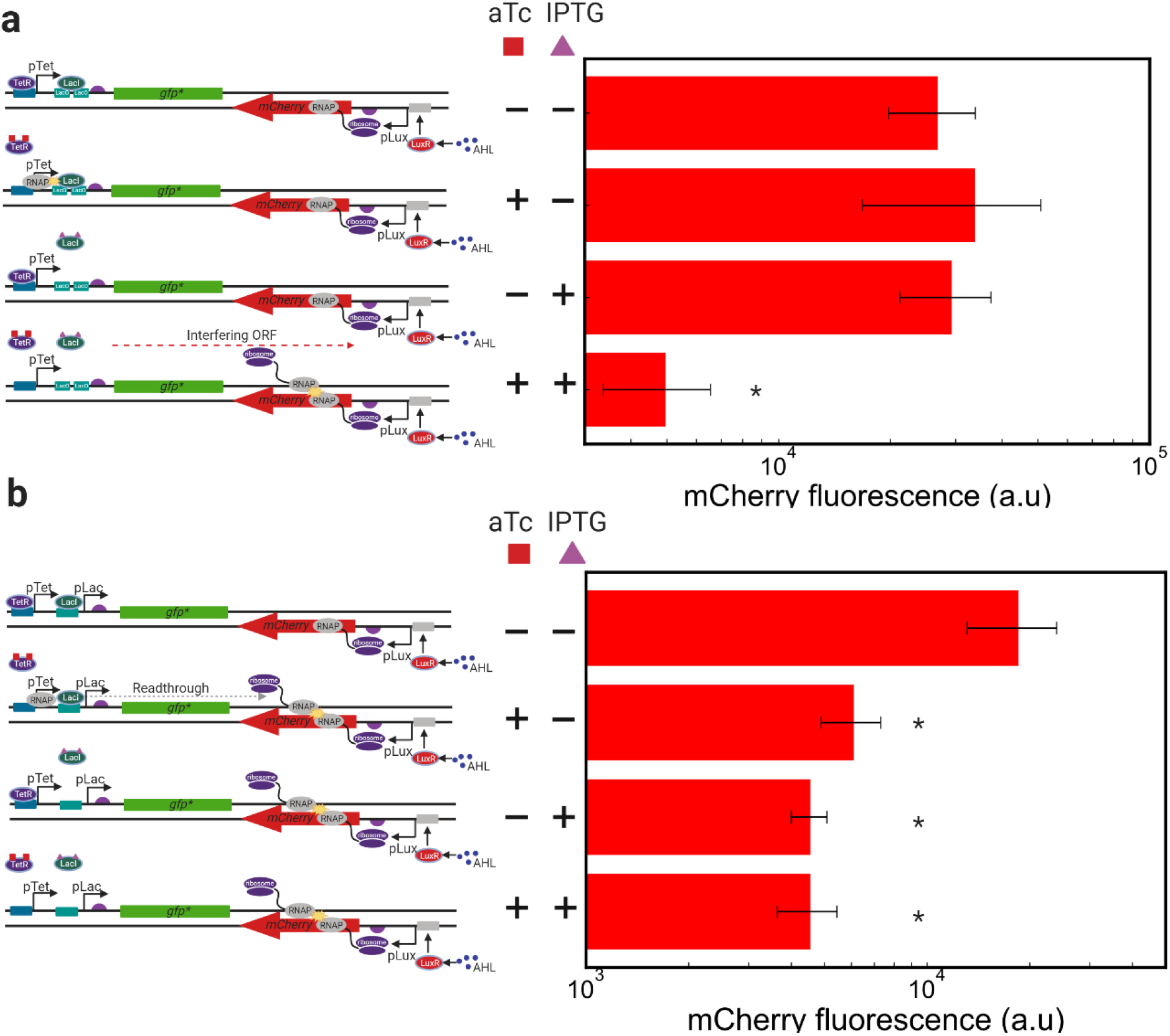
NAND and NOR behaviors arise from coupled roadblock and collisions. Inverting the orientation of a gene of interest in an AND or OR gate creates NAND and NOR logic via TI collisions. a) Using AND logic with an inducible pTet promoter and LacO operator to control the release of interfering expressomes to collide an interfere mCherry expression creates NAND logic behavior. mCherry expression is plotted with i) no inducer, ii) saturating aTc only, iii) saturating IPTG only, and iv) satuarting aTc + IPTG, revealing NAND behavior for this construct. b) Using OR logic with a tandem pTet and pLac promoter system generates NOR logic behavior. mCherry expression is plotted with i) no inducer, ii) saturating aTc only, iii) saturating IPTG only, and iv) satuarting aTc + IPTG, revealing NOR behavior for this construct. * denotes statistical significance compared to the ‘ON’ condition (*p*<0.05, Mann-Whitney *U* test, with n>3 replicates.)

To create NOR behavior, we used a tandem promoter system composed of pLac and pTet, which was previously shown to demonstrate OR behavior (34) to the *gfp*-mCherry* system to produce strong collisions that repress mCherry expression (Fig. 4b, left). Note that the binding affinity of LacI to the LacO binding sites in the downstream promoter was weakened through point mutations in the LacO sequence in order to increase readthrough of RNAP from the upstream promoter(34). We observed a significant ~4-fold decrease in mCherry expression when either aTc, IPTG, or both were present (Fig. 4b, left) at relatively low AHL concentrations (20 μL). We note that at higher AHL concentrations, fold-change due to TI increases, but the NOR behavior grows asymmetric (34, 39), as the induction of both pTet and pLac at saturating conditions represses mCherry expression further than when either was individually activated (Supplementary Figure S9). This additive effect of the tandem promoters could be due to increased interfering RNAP firing, cooperative readthrough of the tandem RNAPs (40), and/or reduced promoter clogging (34, 41). Together, these results demonstrate the first use of TI collisions for the engineering of higher-order genetic devices.

## DISCUSSION

The role of TI in genome-wide regulation and genome organization is still being uncovered. Recently, bacterial transcriptome studies have provided a high resolution picture of the *E. coli* transcriptome and revealed a close relationship between factor-dependent transcriptional termination and TI (5, 7, 26). Here we applied these lessons toward the design of synthetic constructs in which the processivity of an interfering RNAP is engineered to improve the strength of gene repression through transcriptional collisions. We employed three processivity control strategies—the use of the Rho-inhibiting antibiotic bicyclomycin (Fig. 1b-c), the phage polarity suppressing protein Psu (Fig. 1d-f), and the co-translation with the ribosome (Fig. 2)—to tune the strength of TI collisions. We demonstrated, on a transcription-level, the improved processivity control when expressomes are permitted to enter a convergently oriented ORF (Fig. 2d) and the resulting ~38-fold reduction in transcript due to TI (Fig. 2e). We show that changing the expression level of Psu (Fig. 1f), changing strengths of interfering and expressing promoters (Supplementary Figure S7), and adjusting the length with which the expressome can interfere (Fig. 3b) can further tune TI and provide mechanistic insights into the role of antitermination in strengthening TI. We then coupled two modes of TI—roadblock and collisions—to create two-input minimal NAND and NOR logic gates (Fig. 4a-b), representing the first functionally complete Boolean gates constructed using TI collisions. Taken together, these results underscore the importance of RNAP processivity to TI and also demonstrate the tunability of processivity control to engineering TI-based genetic responses.

Moreover, these results further emphasize the close connection between RNAP processivity and TI in the genome. Bacteria manipulate RNAP processivity through several different antitermination mechanisms (42) including RNA aptamers (25), which were recently found to curb levels of antisense transcription and TI throughout the *E. Coli* genome (26). This suggests an evolved strategy to avoid potentially harmful TI. Conversely, the recent discovery that TI is utilized as a widespread bidirectional terminator in *E. coli* (5) raises the interesting prospect that ostensibly destructive collisions between RNAPs are evolutionarily selected for, and that genomes are in some part organized to utilize RNAP collisions for gene regulation in a small genetic space. A recent report demonstrated that head-on collisions of replisome and RNAPs increase the evolvability of convergently-oriented genes through mutagenesis (43), suggesting an evolutionary selection for these convergent arrangements. Collisions between RNAPs, too, could be useful under certain circumstances. It was recently observed that a ‘non-contiguous operon’ governing menaquinone synthesis in *S. aureus* uses antisense transcription to selectively downregulate gene expression to express drug-tolerant small-colony phenotypes (44). These findings have stirred interest in TI as a mechanism shaping evolution. Extending the results of this study to the bacterial genome, it seems that the cell’s ability to alter the processivity of an RNAP through Rho-dependent transcriptional termination suggests that TI in the genome is potentially ‘tunable’. Indeed, several laboratory adaptation studies for different organisms under different stresses (45–47) have found common Rho and RNAP mutations, suggesting a potential role for TI in bacterial stress responses.

If genomes are arranged to facilitate collisions for regulation, could synthetic circuits also take on such an organization? Here we sought to expand TI’s potential for building genetic devices by engineering processive interfering RNAPs and introducing three distinct methods for processivity control. We note that some synthetic biology applications may require gene knockdowns higher than the 38-fold and 10-fold changes in transcript and protein, respectively, reported here. TI systems are capable of ~100-fold changes in gene expression (19, 38), but performance has been shown to depend on gene architecture, promoter strengths, and terminator strengths (9, 18–21). Optimization of these parts, in concert with the processivity control strategy detailed here, should further expand TI’s potential for synthetic biology. Recently, Krylov and colleagues demonstrated the use of an ‘actuator’ sequence element consisting of an antisense promoter, antitermination sequence to protect the antisense RNAPs from Rho termination, and an RNAse III processing sequence used to downregulate expression of three *E. Coli* metabolism genes (38). The application of processivity control for metabolic engineering further demonstrates the applicability of this strategy to synthetic biology.

Despite notable recent works, TI is still largely understudied, and the ‘rules’ determining where, when, and how RNAPs and/or the expressomes collide are not well understood. Rates of transcription and translation both in the genome and on plasmids are highly context-dependent (17, 48, 49) and depend on the position in an operon and proximity to other genes or genetic elements. Moreover, the role of supercoiling in mediating TI collisions is not well understood but is likely important in determining the strength and location of RNAP collisions in genomes across the kingdoms of life. Fundamental insights into these ‘road rules’ (41) of RNAP traffic on the DNA and the resulting TI will reveal how these molecular transcriptional events shape cell physiology and evolution.

## SUPPLEMENTARY DATA

Supplementary Data are available at NAR Online.

## ACKNOWLEDGEMENTS

The authors wish to acknowledge the GAANN fellowship given to N.J.O. through the Department of Education, the S10ODO21601 Grant given to the Flow Cytometry Facility of the University of Colorado Boulder, the Next-Gen Sequencing Core at the University of Colorado Boulder, and the National Science Foundation Grant No. MCB1714564 to A.C.

## AUTHOR CONTRIBUTIONS

N.J.O, A.E.B, and A.C designed the study. N.J.O performed the experiments and analyzed the data. N.J.O and A.C wrote the paper.

