## Supplementary Information for "Engineering Transcriptional Interference through RNA Polymerase Processivity Control"

### Table of Contents


| | |
|---|---|
| Page 2 | Supplemental Note 1: asRNA hybridization in the gfp*-mCherry system |
| Page 4 | Supplementary Table S1: Plasmid backbone architecture and DNA sequences. |
| Page 6 | Supplementary Table S2: Plasmid insert architecture and DNA sequences. |
| Page 8 | Supplementary Table S3: mCherry sequences mutated to introduce antisense stop codons. |
| Page 10 | Supplementary Figure S1: Maps of plasmids used in this study |
| Page 11 | Supplementary Figure S2: Convergent promoters without processivity control produces weak TI. |
| Page 12 | Supplementary Figure S3: High Psu expression increases gene expression under all conditions |
| Page 13 | Supplementary Figure S4: Psu expression exhibits negligible growth effects.. |
| Page 14 | Supplementary Figure S5: Reduction in GFP expression due to TI is observed in the *gfp-mCherry convergent construct.* |
| Page 15 | Supplementary Figure S6: Truncated mCherry transcripts are present in the presence and absence of TI. |
| Page 16 | Supplementary Figure S7: Expression control with processivity control yields a layered TI response. |
| Page 17 | Supplementary Figure S8: TI fold-change over the *gfp*-mCherry* is not a function of expressing promoter strength (AHL) when concentrations exceed 20 µL. |
| Page 18 | Supplementary Figure S9: NOR behavior changes with expressing promoter strength |
| Page 19 | References |

**Supplemental Note 1:**

TI is composed primarily of interactions between transcriptional machinery and interactions between overlapping transcripts. Attempts to parse the contributions of these two mechanisms have revealed either partial[1,2] or negligible[3] contributions of asRNA hybridization. We believe that here, in our system—specifically in the expressome-on-expressome collision system composed of a convergent *gfp** and *mCherry* (Fig. 2a, middle)—RNAP collisions are the dominant form of interference.

The insertion of a unidirectional terminator between the *gfp** and *mCherry* sequences on the *gfp** strand prevents expressomes from reading into the *mCherry* ORF. The lack of a transcriptional terminator on the *mCherry* strand likely allows RNAPs originating from pLux to transcribe into the antisense *gfp** prior to factor-dependent termination. It is possible, then, to have hybridization of sense and antisense RNA *gfp** transcripts that would result in RNase degradation and reduce mCherry expression. However, no reduction in mCherry observed when pTet-LacO is activated (Fig. 3b, bottom) in the *gfp*-mCherry* convergent construct containing a unidirectional terminator, indicating negligible asRNA interference.

Moreover, the observed dependence on TI strength with the length of the interfering ORF (Fig. 4c) also indicates RNAP collisions as the dominant TI mechanism. If interfering ORF length were roughly indicative of antisense transcript length (RNAPs after decoupling may experience factor-dependent termination (Fig. 2c)), the observed trend could be interpreted as longer antisense transcripts with more overlap in the *mCherry* sequence causing a reduction in mCherry expression due to hybridization. However, hybridization of asRNAs would not likely exhibit such a transcript length dependence due to the formation of RNA secondary structures that may reduce hybridization affinity. Rather, the observed trend in TI with interfering ORF length is more consistent with a change in collisions ‘location’ on the DNA: RNAPs that remain coupled with ribosomes far into the *mCherry* ORF are more likely to, on average, result in TI prior to the complete transcription of *mCherry*, resulting here in a stronger TI fold-change. These two experiments demonstrate that TI collisions are the dominant mode of gene repression in our system instead of mRNA hybridization.

**Supplementary Table S1**: **Plasmid backbone architecture and DNA sequences**. The two listed plasmid backbones were used for all constructs in this study. Abbreviations are as follows: LuxR- transcriptional activator of pLux; BBa23108- constitutive promoter; ColE1 ori- high copy number origin of replication; KanR – Kanamycin resistance cassette; Psu- polarity suppression gene from bacteriophage P4; araC- arabinose regulatory protein, pBad activator; pBad- arabinose-inducible promoter. Underlined features are oriented on the opposite strand relative to pTet

| Plasmid Architecture | Sequence |
|---|---|
| rrnBT1 terminator—LuxR—Bba23108—ColE1 ori – lambda t0 terminator-- KanR | ACCGGTCATGTTCTTTCCTGCGTTATCCCCTGATTCTGTGGATAACCGTATTACCGCCTTTGAGTGAGCTG<br>ATACCGCTCGCCGCAGCCGAACGCCATAGGTCTAGGGCGGCGGATTTGTCCTACTCAGGAGAGCGTTCAC<br>CGACAAACAACAGATAAAACGAAAGGCCCAGTCTTTCGACTGAGCCTTTCGTTTTATTTGGGGCCCTTAA<br>TTTTTAAAGTATGGGCAATCAATTGCTCCTGTTAAAATTGCTTTAGAAATACTTTGGCAGCGGTTTGTTGT<br>ATTGAGTTTCATTTGCGCATTGGTTAAATGGAAAGTGACAGTACGCTCACTGCAGCCTAATATTTTTGAAA<br>TATCCCAAGAGCTTTTTCCTTCGCATGCCCACGCTAAACATTCTTTTTCTCTTTTGGTTAAATCGTTGTTTG<br>ATTTATTATTTGCTATATTTATTTTTCGATAATTATCAACTAGAGAAGGAACAATTAATGGTATGTTCATA<br>CACGCATGTAAAAATAAACTATCTATATAGTTGTCTTTTTCTGAATGTGCAAAACTAAGCATTCCGAAGCC<br>ATTGTTAGCCGTATGAATAGGGAAACTAAACCCAGTGATAAGACCTGATGTTTTCGCTTCTTTAATTACAT<br>TTGGAGATTTTTTATTTACAGCATTGTTTTCAAATATATTCCAATTAATTGGTGAATGATTGGAGTTAGAA<br>TAATCTACTATAGGATCATATTTTATTAAATTAGCGTCATCATAATATTGCCTCCATTTTTTAGGGTAATTA<br>TCTAGAATTGAAATATCAGATTTAACCATAGAATGAGGATAAATGATCGCGAGTAAATAATATTCACAAT<br>GTACCATTTTAGTCATATCAGATAAGCATTGATTAATATCATTATTGCTTCTACAAGCTTTAATTTTATTAA<br>TTATTCTGTATGTGTCGTCGGCATTTATGTTTTTCATACCCATCTCTTTATCCTTACCTATTGTTTGTCGCAA<br>GTTTTGCGTGTTATATATCATTAAAACGGTAATGGATTGACATTTGATTCTAATAAATTAGATGCATGGGC<br>CCGCTAGCATTATACCTAGGACTGAGCTAGCTGTCAGACATGTGAGCAAAAGGCCAGCAAAAGGCCAGG<br>AACCGTAAAAAGGCCGCGTTGCTGGCGTTTTTCCATAGGCTCCGCCCCCCTGACGAGCATCACAAAAATC<br>GACGCTCAAGTCAGAGGTGGCGAAACCCGACAGGACTATAAAGATACCAGGCGTTTCCCCCTGGAAGCT<br>CCCTCGTGCGCTCTCCTGTTCCGACCCTGCCGCTTACCGGATACCTGTCCGCCTTTCTCCCTTCGGGAAGC<br>GTGGCGCTTTCTCAATGCTCACGCTGTAGGTATCTCAGTTCGGTGTAGGTCGTTCGCTCCAAGCTGGGCTG<br>TGTGCACGAACCCCCCGTTCAGCCCGACCGCTGCGCCTTATCCGGTAACTATCGTCTTGAGTCCAACCCGG<br>TAAGACACGACTTATCGCCACTGGCAGCAGCCACTGGTAACAGGATTAGCAGAGCGAGGTATGTAGGCG<br>GTGCTACAGAGTTCTTGAAGTGGTGGCCTAACTACGGCTACACTAGAAGGACAGTATTTGGTATCTGCGC<br>TCTGCTGAAGCCAGTTACCTTCGGAAAAAGAGTTGGTAGCTCTTGATCCGGCAAACAAACCACCGCTGGT<br>AGCGGTGGTTTTTTTGTTTGCAAGCAGCAGATTACGCGCAGAAAAAAAGGATCTCAAGAAGATCCTTTGA<br>TCTTTTCTACGGGGTCTGACGCTCAGTGGAACGAAAACTCACGTTAAGGGATTTTGGTCATGACTAGTGCT<br>TGGATTCTCACCAATAAAAAACGCCCGGCGGCAACCGAGCGTTCTGAACAAATCCAGATGGAGTTCTGAG<br>GTCATTACTGGATCTATCAACAGGAGTCCAAGCGAGCTCTCGAACCCCAGAGTCCCGCTCAGAAGAACTC<br>GTCAAGAAGGCGATAGAAGGCGATGCGCTGCGAATCGGGAGCGGCGATACCGTAAAGCACGAGGAAGC<br>GGTCAGCCCATTCGCCGCCAAGCTCTTCAGCAATATCACGGGTAGCCAACGCTATGTCCTGATAGCGGTC<br>CGCCACACCCAGCCGGCCACAGTCGATGAATCCAGAAAAGCGGCCATTTTCCACCATGATATTCGGCAAG<br>CAGGCATCGCCATGGGTCACGACGAGATCCTCGCCGTCGGGCATGCGCGCCTTGAGCCTGGCGAACAGTT<br>CGGCTGGCGCGAGCCCCTGATGCTCTTCGTCCAGATCATCCTGATCGACAAGACCGGCTTCCATCCGAGT<br>ACGTGCTCGCTCGATGCGATGTTTCGCTTGGTGGTCGAATGGGCAGGTAGCCGGATCAAGCGTATGCAGC<br>CGCCGCATTGCATCAGCCATGATGGATACTTTCTCGGCAGGAGCAAGGTGAGATGACAGGAGATCCTGCC<br>CCGGCACTTCGCCCAATAGCAGCCAGTCCCTTCCCGCTTCAGTGACAACGTCGAGCACAGCTGCGCAAGG<br>AACGCCCGTCGTGGCCAGCCACGATAGCCGCGCTGCCTCGTCCTGCAGTTCATTCAGGGCACCGGACAGG<br>TCGGTCTTGACAAAAAGAACCGGGCGCCCCTGCGCTGACAGCCGGAACACGGCGGCATCAGAGCAGCCG<br>ATTGTCTGTTGTGCCCAGTCATAGCCGAATAGCCTCTCCACCCAAGCGGCCGGAGAACCTGCGTGCAATC<br>CATCTTGTTCAATCATGCGAAACGATCCTCATCCTGTCTCTTGATCAGATCTTGATCCCCTGCGCCATCAG<br>ATCCTTGGCGGCAAGAAAGCCATCCAGTTTACTTTGCAGGGCTTCCCAACCTTACCAGAGGGCGCCCCAG<br>CTGGCAATTCCGACGTCTAAGAAACCATTATTATCATGACATTAACCTATAAAAATAGGCGTATCACGAG<br>GCCCTTTCGTCTTCACCTCGAG |

| | |
|---|---|
| rrnBT1 terminator—LuxR—Bba23108—Psu—pBad—araC*—tonb terminator –ColE1 ori – lambda t0 terminator--KanR | ACCGGTCATGTTCTTTCCTGCGTTATCCCCTGATTCTGTGGATAACCGTATTACCGCCTTTGAGTGAGCTG<br>ATACCGCTCGCCGCAGCCGAACGCCATAGGTCTAGGGCGGCGGATTTGTCCTACTCAGGAGAGCGTTCAC<br>CGACAAACAACAGATAAAACGAAAGGCCCAGTCTTTCGACTGAGCCTTTCGTTTTATTTGGGGCCCTTAA<br>TTTTTAAAGTATGGGCAATCAATTGCTCCTGTTAAAATTGCTTTAGAAATACTTTGGCAGCGGTTTGTTGT<br>ATTGAGTTTCATTTGCGCATTGGTTAAATGGAAAGTGACAGTACGCTCACTGCAGCCTAATATTTTTGAAA<br>TATCCCAAGAGCTTTTTCCTTCGCATGCCCACGCTAAACATTCTTTTTCTCTTTTGGTTAAATCGTTGTTTG<br>ATTTATTATTTGCTATATTTATTTTTCGATAATTATCAACTAGAGAAGGAACAATTAATGGTATGTTCATA<br>CACGCATGTAAAAATAAACTATCTATATAGTTGTCTTTTTCTGAATGTGCAAAACTAAGCATTCCGAAGCC<br>ATTGTTAGCCGTATGAATAGGGAAACTAAACCCAGTGATAAGACCTGATGTTTTCGCTTCTTTAATTACAT<br>TTGGAGATTTTTTATTTACAGCATTGTTTTCAAATATATTCCAATTAATTGGTGAATGATTGGAGTTAGAA<br>TAATCTACTATAGGATCATATTTTATTAAATTAGCGTCATCATAATATTGCCTCCATTTTTTAGGGTAATTA<br>TCTAGAATTGAAATATCAGATTTAACCATAGAATGAGGATAAATGATCGCGAGTAAATAATATTCACAAT<br>GTACCATTTTAGTCATATCAGATAAGCATTGATTAATATCATTATTGCTTCTACAAGCTTTAATTTTATTAA<br>TTATTCTGTATGTGTCGTCGGCATTTATGTTTTTCATACCCATCTCTTTATCCTTACCTATTGTTTGTCGCAA<br>GTTTTGCGTGTTATATATCATTAAAACGGTAATGGATTGACATTTGATTCTAATAAATTAGATGCATGGGC<br>CCGCTAGCATTATACCTAGGACTGAGCTAGCTGTCAACATGTcgcgtaccatgggcccTtacactgactgacgtgatgccagttgcg<br>cacttttacgcgaaaaaatcatgttctgcgcaggggtgaatttttcacggctgtcatccaccgaagccacgtcaggcctgaatccgatggtcgttaaaatgtcgctatcctgtgcg<br>gaataattaatttttttcacccttcgcaagccaggacagaagggcttcacgcagggcatctgtggcacgctgtatggcacagtttcgggcaatggccgtcagctcactgtagccc<br>atcagctccggtgccagtgcggccgccagtgctgtgccgtgctgctgcataaaatcattcagccggtcgcggatgctgatgtgctgaacggcttcatgcgaacgaatataacg<br>accggcggcctgattcacctgccattttctgacttcgataatattgcgtaattcgtccaggcgactgacgtttctgccttctcctgacagaagccgcagatattcctgttcggccgct<br>gccagctcatttttgcgttgcagccatgctgctttgttattctgacaggtgtcaaaggcctgctgtaaggctgtgctttccatggtatatCTCCTTcttaaagttcgtatctccgcc<br>tatctcacacgagataggcggtgctagcccaaaaaaacgggtatggagaaacagtagagagttgcgataaaaagcgtcaggtaggatccgctaatcttatggataaaaatgct<br>atggcatagcaaagtgtgacgccgtgcaaataatcaatgtggacttttctgccgtgattatagacacttttgttacgcgtttttgtcatggctttggtcccgctttgttacagaatgcttt<br>taataagcggggttaccggtttggttagcgagaagagccagtaaaagacgcagtgacggcaatgtctgatgcaatatggacaattggtttcttctctgaatggcgggagtatga<br>aaagtatggctgaagcgcaaaatgatcccctgctgccgggatactcgtttaatgcccatctggtggcgggtttaacgccgattgaggccaacggttatctcgatttttttatcgacc<br>gaccgctgggaatgaaaggttatattctcaatctcaccattcgcggtcagggggtggtgaaaaatcagggacgagaatttgtttgccgaccgggtgatattttgctgttcccgcc<br>aggagagattcatcactacggtcgtcatccggaggctcgcgaatggtatcaccagtgggtttactttcgtccgcgcgcctactggcatgaatggcttaactggccgtcaatatttg<br>ccaatacggggttctttcgcccggatgaagcgcaccagccgcatttcagcgacctgtttgggcaaatcattaacgccgggcaaggggaagggcgctattcggagctgctggc<br>gataaatctgcttgagcaattgttactgcggcgcatggaagcgattaacgagtcgctccatccaccgatggataatcgggtacgcgaggcttgtcagtacatcagcgatcacct<br>ggcagacagcaattttgatatcgccagcgtcgcacagcatgtttgcttgtcgccgtcgcgtctgtcacatcttttccgccagcagttagggattagcgtcttaagctggcgcgag<br>gaccaacgtatcagccaggcgaagctgcttttgagcaccacccggatgcctatcgccaccgtcggtcgcaatgttggttttgacgatcaactctatttctcgcgggtatttaaaaa<br>atgcaccggggccagcccgagcgagttccgtgccggttgagaagaaaaagtgaatgatgtagccgtcaagttgtcataaaaatcatacctgacctccatagcagaaagtcaa<br>aagcctccgaccggaggcttttgactGACATGTGAGCAAAAGGCCAGCAAAAGGCCAGGAACCGTAAAAAGGCCGCGTT<br>GCTGGCGTTTTTCCATAGGCTCCGCCCCCCTGACGAGCATCACAAAAATCGACGCTCAAGTCAGAGGTGG<br>CGAAACCCGACAGGACTATAAAGATACCAGGCGTTTCCCCCTGGAAGCTCCCTCGTGCGCTCTCCTGTTC<br>CGACCCTGCCGCTTACCGGATACCTGTCCGCCTTTCTCCCTTCGGGAAGCGTGGCGCTTTCTCAATGCTCA<br>CGCTGTAGGTATCTCAGTTCGGTGTAGGTCGTTCGCTCCAAGCTGGGCTGTGTGCACGAACCCCCCGTTCA<br>GCCCGACCGCTGCGCCTTATCCGGTAACTATCGTCTTGAGTCCAACCCGGTAAGACACGACTTATCGCCA<br>CTGGCAGCAGCCACTGGTAACAGGATTAGCAGAGCGAGGTATGTAGGCGGTGCTACAGAGTTCTTGAAG<br>TGGTGGCCTAACTACGGCTACACTAGAAGGACAGTATTTGGTATCTGCGCTCTGCTGAAGCCAGTTACCTT<br>CGGAAAAAGAGTTGGTAGCTCTTGATCCGGCAAACAAACCACCGCTGGTAGCGGTGGTTTTTTTGTTTGC<br>AAGCAGCAGATTACGCGCAGAAAAAAAGGATCTCAAGAAGATCCTTTGATCTTTTCTACGGGGTCTGACG<br>CTCAGTGGAACGAAAACTCACGTTAAGGGATTTTGGTCATGACTAGTGCTTGGATTCTCACCAATAAAAA<br>ACGCCCGGCGGCAACCGAGCGTTCTGAACAAATCCAGATGGAGTTCTGAGGTCATTACTGGATCTATCAA<br>CAGGAGTCCAAGCGAGCTCTCGAACCCCAGAGTCCCGCTCAGAAGAACTCGTCAAGAAGGCGATAGAAG<br>GCGATGCGCTGCGAATCGGGAGCGGCGATACCGTAAAGCACGAGGAAGCGGTCAGCCCATTCGCCGCCA<br>AGCTCTTCAGCAATATCACGGGTAGCCAACGCTATGTCCTGATAGCGGTCCGCCACACCCAGCCGGCCAC<br>AGTCGATGAATCCAGAAAAGCGGCCATTTTCCACCATGATATTCGGCAAGCAGGCATCGCCATGGGTCAC<br>GACGAGATCCTCGCCGTCGGGCATGCGCGCCTTGAGCCTGGCGAACAGTTCGGCTGGCGCGAGCCCCTGA<br>TGCTCTTCGTCCAGATCATCCTGATCGACAAGACCGGCTTCCATCCGAGTACGTGCTCGCTCGATGCGATG<br>TTTCGCTTGGTGGTCGAATGGGCAGGTAGCCGGATCAAGCGTATGCAGCCGCCGCATTGCATCAGCCATG<br>ATGGATACTTTCTCGGCAGGAGCAAGGTGAGATGACAGGAGATCCTGCCCCGGCACTTCGCCCAATAGCA<br>GCCAGTCCCTTCCCGCTTCAGTGACAACGTCGAGCACAGCTGCGCAAGGAACGCCCGTCGTGGCCAGCCA<br>CGATAGCCGCGCTGCCTCGTCCTGCAGTTCATTCAGGGCACCGGACAGGTCGGTCTTGACAAAAAGAACC<br>GGGCGCCCCTGCGCTGACAGCCGGAACACGGCGGCATCAGAGCAGCCGATTGTCTGTTGTGCCCAGTCAT<br>AGCCGAATAGCCTCTCCACCCAAGCGGCCGGAGAACCTGCGTGCAATCCATCTTGTTCAATCATGCGAAA<br>CGATCCTCATCCTGTCTCTTGATCAGATCTTGATCCCCTGCGCCATCAGATCCTTGGCGGCAAGAAAGCCA<br>TCCAGTTTACTTTGCAGGGCTTCCCAACCTTACCAGAGGGCGCCCCAGCTGGCAATTCCGACGTCTAAGA<br>AACCATTATTATCATGACATTAACCTATAAAAATAGGCGTATCACGAGGCCCTTTCGTCTTCACCTCGAG |

.

**Supplementary Table S2**: **Plasmid insert architecture and DNA sequences**. The listed plasmid inserts were combined with one or both of the plasmid backbones in Supplementary Table S1 to form the constructs used in this study. Abbreviations are as follows: mCherry – red fluorescent protein; gfp- green fluorescent protein; gfp*- green fluorescent protein sequence without a stop codon; rrnBT1- terminator sequence; pTet+LacO- a tandem TetR-repressable promoter, pTet upstream two repressible Lac operator sequences[4]. BioBrick suffix was obtained from Twist Biosciences. All underlined features are oriented antisense to pTet.

| | |
|---|---|
| pTet+LacO – *gfp* – BioBrick Suffix - pLux | 5’ -<br>TCCCTATCAGTGATAGAGATTGACATCCCTATCAGTGATAGAGATACTGAGCACATCAGCAGGACGCACTGACC<br>GAATTCATTAAAGAGGAGAAAGGTACCTTGTGAGCGGATAACAAAAAGGCTTGTGAGCGGATAACAAACCTGG<br>GTCGACAATCACGCGTACCATGGGATCCCTATTTGTATAGTTCATCCATGCCATGTGTAATCCCAGCAGCTGTTA<br>CAAACTCAAGAAGGACCATGTGGTCTCTCTTTTCGTTGGGATCTTTCGAAAGGGCAGATTGTGTGGACAGGTAA<br>TGGTTGTCTGGTAAAAGGACAGGGCCATCGCCAATTGGAGTATTTTGTTGATAATGGTCTGCTAGTTGAACGCTT<br>CCATCTTCAATGTTGTGTCTAATTTTGAAGTTAACTTTGATTCCATTCTTTTGTTTGTCTGCCATGATGTATACATT<br>GTGTGAGTTATAGTTGTATTCCAATTTGTGTCCAAGAATGTTTCCATCTTCTTTAAAATCAATACCTTTTAACTCG<br>ATTCTATTAACAAGGGTATCACCTTCAAACTTGACTTCAGCACGTGTCTTGTAGTTCCCGTCATCTTTGAAAAAT<br>ATAGTTCTTTCCTGTACATAACCTTCGGGCATGGCACTCTTGAAAAAGTCATGCCGTTTCATATGATCTGGGTAT<br>CTTGAAAAGCATTGAACACCATAAGAGAAAGTAGTGACAAGTGTTGGCCATGGAACAGGTAGTTTTCCAGTAGT<br>GCAAATAAATTTAAGGGTAAGTTTTCCGTATGTTGCATCACCTTCACCCTCTCCACTGACAGAAAATTTGTGCCC<br>ATTAACATCACCATCTAATTCAACAAGAATTGGGACAACTCCAGTGAAAAGTTCTTCTCCTTTACTCATCCTTCT<br>CTCCTCTGCAGCGGCCGCTACTAGTATTCATTCGACTATAACAAACCATTTTCTTGCGTAAACCTGTACGATCCT<br>ACAGGT – 3’ |
| pTet+LacO – *gfp** – *mCherry*— BioBrick Suffix - pLux | 5’ -<br>TCCCTATCAGTGATAGAGATTGACATCCCTATCAGTGATAGAGATACTGAGCACATCAGCAGGACGCACTGACC<br>GAATTCATTAAAGAGGAGAAAGGTACCTTGTGAGCGGATAACAAAAAGGCTTGTGAGCGGATAACAAACCTGG<br>GTCGACAAGGAGAGAAGGATGAGTAAAGGAGAAGAACTTTTCACTGGAGTTGTCCCAATTCTTGTTGAATTAGA<br>TGGTGATGTTAATGGGCACAAATTTTCTGTCAGTGGAGAGGGTGAAGGTGATGCAACATACGGAAAACTTACCC<br>TTAAATTTATTTGCACTACTGGAAAACTACCTGTTCCATGGCCAACACTTGTCACTACTTTCTCTTATGGTGTTCA<br>ATGCTTTTCAAGATACCCAGATCATATGAAACGGCATGACTTTTTCAAGAGTGCCATGCCCGAAGGTTATGTAC<br>AGGAAAGAACTATATTTTTCAAAGATGACGGGAACTACAAGACACGTGCTGAAGTCAAGTTTGAAGGTGATAC<br>CCTTGTTAATAGAATCGAGTTAAAAGGTATTGATTTTAAAGAAGATGGAAACATTCTTGGACACAAATTGGAAT<br>ACAACTATAACTCACACAATGTATACATCATGGCAGACAAACAAAAGAATGGAATCAAAGTTAACTTCAAAATT<br>AGACACAACATTGAAGATGGAAGCGTTCAACTAGCAGACCATTATCAACAAAATACTCCAATTGGCGATGGCCC<br>TGTCCTTTTACCAGACAACCATTACCTGTCCACACAATCTGCCCTTTCGAAAGATCCCAACGAAAAGAGAGACC<br>ACATGGTCCTTCTTGAGTTTGTAACAGCTGCTGGGATTACACATGGCATGGATGAACTATACAAACAGGGATCC<br>TTACTTGTACAGCTCGTCCATGCCGCCGGTGGAGTGGCGGCCCTCGGCGCGTTCGTACTGTTCCACGATGGTGTA<br>GTCCTCGTTGTGGGAGGTGATGTCCAACTTGATGTTGACGTTGTAGGCGCCGGGCAGCTGCACGGGCTTCTTGGC<br>CTTGTAGGTGGTCTTGACCTCAGCGTCGTAGTGGCCGCCGTCCTTCAGCTTCAGCCTCTGCTTGATCTCGCCCTTC<br>AGGGCGCCGTCCTCGGGGTACATCCGCTCGGAGGAGGCCTCCCAGCCCATGGTCTTCTTCTGCATTACGGGGCC<br>GTCGGAGGGGAAGTTGGTGCCGCGCAGCTTCACCTTGTAGATGAACTCGCCGTCCTGCAGGGAGGAGTCCTGGG<br>TCACGGTCACCACGCCGCCGTCCTCGAAGTTCATCACGCGCTCCCACTTGAAGCCCTCGGGGAAGGACAGCTTC<br>AAGTAGTCGGGGATGTCGGCGGGGTGCTTCACGTAGGCCTTGGAGCCGTACATGAACTGAGGGGACAGGATGT<br>CCCAGGCGAAGGGCAGGGGGCCACCCTTGGTCACCTTCAGCTTGGCGGTCTGGGTGCCCTCGTAGGGGCGGCCC<br>TCGCCCTCGCCCTCGATCTCGAACTCGTGGCCGTTCACGGAGCCCTCCATGTGCACCTTGAAGCGCATGAACTCC<br>TTGATGATGGCCATGTTATCCTCCTCGCCCTTGCTCACCAT CCTTCTCTCCT<br>CTGCAGCGGCCGCTACTAGTATTCATTCGACTATAACAAACCATTTTCTTGCGTAAACCTGTACGATCCTACAGG<br>T – 3’ |
| pTet+LacO – *gfp*– *mCherry*— BioBrick Suffix - pLux | 5’ -<br>TCCCTATCAGTGATAGAGATTGACATCCCTATCAGTGATAGAGATACTGAGCACATCAGCAGGACGCACTGACC<br>GAATTCATTAAAGAGGAGAAAGGTACCTTGTGAGCGGATAACAAAAAGGCTTGTGAGCGGATAACAAACCTGG<br>GTCGACAAGGAGAGAAGG<br>ATGAGTAAAGGAGAAGAACTTTTCACTGGAGTTGTCCCAATTCTTGTTGAATTAGATGGTGATGTTAATGGGCA<br>CAAATTTTCTGTCAGTGGAGAGGGTGAAGGTGATGCAACATACGGAAAACTTACCCTTAAATTTATTTGCACTA<br>CTGGAAAACTACCTGTTCCATGGCCAACACTTGTCACTACTTTCTCTTATGGTGTTCAATGCTTTTCAAGATACCC<br>AGATCATATGAAACGGCATGACTTTTTCAAGAGTGCCATGCCCGAAGGTTATGTACAGGAAAGAACTATATTTT<br>TCAAAGATGACGGGAACTACAAGACACGTGCTGAAGTCAAGTTTGAAGGTGATACCCTTGTTAATAGAATCGAG<br>TTAAAAGGTATTGATTTTAAAGAAGATGGAAACATTCTTGGACACAAATTGGAATACAACTATAACTCACACAA<br>TGTATACATCATGGCAGACAAACAAAAGAATGGAATCAAAGTTAACTTCAAAATTAGACACAACATTGAAGAT<br>GGAAGCGTTCAACTAGCAGACCATTATCAACAAAATACTCCAATTGGCGATGGCCCTGTCCTTTTACCAGACAA<br>CCATTACCTGTCCACACAATCTGCCCTTTCGAAAGATCCCAACGAAAAGAGAGACCACATGGTCCTTCTTGAGTT<br>TGTAACAGCTGCTGGGATTACACATGGCATGGATGAACTATACAAATAGGGATCCTTACTTGTACAGCTCGTCC<br>ATGCCGCCGGTGGAGTGGCGGCCCTCGGCGCGTTCGTACTGTTCCACGATGGTGTAGTCCTCGTTGTGGGAGGT<br>GATGTCCAACTTGATGTTGACGTTGTAGGCGCCGGGCAGCTGCACGGGCTTCTTGGCCTTGTAGGTGGTCTTGAC<br>CTCAGCGTCGTAGTGGCCGCCGTCCTTCAGCTTCAGCCTCTGCTTGATCTCGCCCTTCAGGGCGCCGTCCTCGGG<br>GTACATCCGCTCGGAGGAGGCCTCCCAGCCCATGGTCTTCTTCTGCATTACGGGGCCGTCGGAGGGGAAGTTGG<br>TGCCGCGCAGCTTCACCTTGTAGATGAACTCGCCGTCCTGCAGGGAGGAGTCCTGGGTCACGGTCACCACGCCG<br>CCGTCCTCGAAGTTCATCACGCGCTCCCACTTGAAGCCCTCGGGGAAGGACAGCTTCAAGTAGTCGGGGATGTC<br>GGCGGGGTGCTTCACGTAGGCCTTGGAGCCGTACATGAACTGAGGGGACAGGATGTCCCAGGCGAAGGGCAGG<br>GGGCCACCCTTGGTCACCTTCAGCTTGGCGGTCTGGGTGCCCTCGTAGGGGCGGCCCTCGCCCTCGCCCTCGATC<br>TCGAACTCGTGGCCGTTCACGGAGCCCTCCATGTGCACCTTGAAGCGCATGAACTCCTTGATGATGGCCATGTTA<br>TCCTCCTCGCCCTTGCTCACCAT CCTTCTCTCCT<br>CTGCAGCGGCCGCTACTAGTATTCATTCGACTATAACAAACCATTTTCTTGCGTAAACCTGTACGATCCTACAGG<br>T – 3’ |

| | |
|---|---|
| pTet+LacO – *mCherry* – BioBrick Suffix - pLux | 5’ -<br>TCCCTATCAGTGATAGAGATTGACATCCCTATCAGTGATAGAGATACTGAGCACATCAGCAGGACGCACTGACC<br>GAATTCATTAAAGAGGAGAAAGGTACCTTGTGAGCGGATAACAAAAAGGCTTGTGAGCGGATAACAAACCTGG<br>GTCGACAATCACGCGTACCATGGGATCC<br>TTACTTGTACAGCTCGTCCATGCCGCCGGTGGAGTGGCGGCCCTCGGCGCGTTCGTACTGTTCCACGATGGTGTA<br>GTCCTCGTTGTGGGAGGTGATGTCCAACTTGATGTTGACGTTGTAGGCGCCGGGCAGCTGCACGGGCTTCTTGGC<br>CTTGTAGGTGGTCTTGACCTCAGCGTCGTAGTGGCCGCCGTCCTTCAGCTTCAGCCTCTGCTTGATCTCGCCCTTC<br>AGGGCGCCGTCCTCGGGGTACATCCGCTCGGAGGAGGCCTCCCAGCCCATGGTCTTCTTCTGCATTACGGGGCC<br>GTCGGAGGGGAAGTTGGTGCCGCGCAGCTTCACCTTGTAGATGAACTCGCCGTCCTGCAGGGAGGAGTCCTGGG<br>TCACGGTCACCACGCCGCCGTCCTCGAAGTTCATCACGCGCTCCCACTTGAAGCCCTCGGGGAAGGACAGCTTC<br>AAGTAGTCGGGGATGTCGGCGGGGTGCTTCACGTAGGCCTTGGAGCCGTACATGAACTGAGGGGACAGGATGT<br>CCCAGGCGAAGGGCAGGGGGCCACCCTTGGTCACCTTCAGCTTGGCGGTCTGGGTGCCCTCGTAGGGGCGGCCC<br>TCGCCCTCGCCCTCGATCTCGAACTCGTGGCCGTTCACGGAGCCCTCCATGTGCACCTTGAAGCGCATGAACTCC<br>TTGATGATGGCCATGTTATCCTCCTCGCCCTTGCTCACCAT CCTTCTCTCCT<br>CTGCAGCGGCCGCTACTAGTATTCATTCGACTATAACAAACCATTTTCTTGCGTAAACCTGTACGATCCTACAGG<br>T – 3’ |
| pTet+LacO w/ RBSMut1– *gfp** –*mCherry*— BioBrick Suffix - pLux | 5’ -<br>TCCCTATCAGTGATAGAGATTGACATCCCTATCAGTGATAGAGATACTGAGCACATCAGCAGGACGCACTGACC<br>GAATTCATTAAAGAGGAGAAAGGTACCTTGTGAGCGGATAACAAAAAGGCTTGTGAGCGGATAACAAACCTGG<br>GTCGACAAGAAGAGAAGG<br>ATGAGTAAAGGAGAAGAACTTTTCACTGGAGTTGTCCCAATTCTTGTTGAATTAGATGGTGATGTTAATGGGCA<br>CAAATTTTCTGTCAGTGGAGAGGGTGAAGGTGATGCAACATACGGAAAACTTACCCTTAAATTTATTTGCACTA<br>CTGGAAAACTACCTGTTCCATGGCCAACACTTGTCACTACTTTCTCTTATGGTGTTCAATGCTTTTCAAGATACCC<br>AGATCATATGAAACGGCATGACTTTTTCAAGAGTGCCATGCCCGAAGGTTATGTACAGGAAAGAACTATATTTT<br>TCAAAGATGACGGGAACTACAAGACACGTGCTGAAGTCAAGTTTGAAGGTGATACCCTTGTTAATAGAATCGAG<br>TTAAAAGGTATTGATTTTAAAGAAGATGGAAACATTCTTGGACACAAATTGGAATACAACTATAACTCACACAA<br>TGTATACATCATGGCAGACAAACAAAAGAATGGAATCAAAGTTAACTTCAAAATTAGACACAACATTGAAGAT<br>GGAAGCGTTCAACTAGCAGACCATTATCAACAAAATACTCCAATTGGCGATGGCCCTGTCCTTTTACCAGACAA<br>CCATTACCTGTCCACACAATCTGCCCTTTCGAAAGATCCCAACGAAAAGAGAGACCACATGGTCCTTCTTGAGTT<br>TGTAACAGCTGCTGGGATTACACATGGCATGGATGAACTATACAAACAGGGATCCTTACTTGTACAGCTCGTCC<br>ATGCCGCCGGTGGAGTGGCGGCCCTCGGCGCGTTCGTACTGTTCCACGATGGTGTAGTCCTCGTTGTGGGAGGT<br>GATGTCCAACTTGATGTTGACGTTGTAGGCGCCGGGCAGCTGCACGGGCTTCTTGGCCTTGTAGGTGGTCTTGAC<br>CTCAGCGTCGTAGTGGCCGCCGTCCTTCAGCTTCAGCCTCTGCTTGATCTCGCCCTTCAGGGCGCCGTCCTCGGG<br>GTACATCCGCTCGGAGGAGGCCTCCCAGCCCATGGTCTTCTTCTGCATTACGGGGCCGTCGGAGGGGAAGTTGG<br>TGCCGCGCAGCTTCACCTTGTAGATGAACTCGCCGTCCTGCAGGGAGGAGTCCTGGGTCACGGTCACCACGCCG<br>CCGTCCTCGAAGTTCATCACGCGCTCCCACTTGAAGCCCTCGGGGAAGGACAGCTTCAAGTAGTCGGGGATGTC<br>GGCGGGGTGCTTCACGTAGGCCTTGGAGCCGTACATGAACTGAGGGGACAGGATGTCCCAGGCGAAGGGCAGG<br>GGGCCACCCTTGGTCACCTTCAGCTTGGCGGTCTGGGTGCCCTCGTAGGGGCGGCCCTCGCCCTCGCCCTCGATC<br>TCGAACTCGTGGCCGTTCACGGAGCCCTCCATGTGCACCTTGAAGCGCATGAACTCCTTGATGATGGCCATGTTA<br>TCCTCCTCGCCCTTGCTCACCAT CCTTCTCTCCT<br>CTGCAGCGGCCGCTACTAGTATTCATTCGACTATAACAAACCATTTTCTTGCGTAAACCTGTACGATCCTACAGG<br>T – 3’ |
| pTet+LacO –*gfp* –BioBrick Suffix - pLux | 5’ -<br>TCCCTATCAGTGATAGAGATTGACATCCCTATCAGTGATAGAGATACTGAGCACATCAGCAGGACGCACTGACC<br>GAATTCATTAAAGAGGAGAAAGGTACCTTGTGAGCGGATAACAAAAAGGCTTGTGAGCGGATAACAAACCTGG<br>GTCGACAAGGAGAGAAGG<br>ATGAGTAAAGGAGAAGAACTTTTCACTGGAGTTGTCCCAATTCTTGTTGAATTAGATGGTGATGTTAATGGGCA<br>CAAATTTTCTGTCAGTGGAGAGGGTGAAGGTGATGCAACATACGGAAAACTTACCCTTAAATTTATTTGCACTA<br>CTGGAAAACTACCTGTTCCATGGCCAACACTTGTCACTACTTTCTCTTATGGTGTTCAATGCTTTTCAAGATACCC<br>AGATCATATGAAACGGCATGACTTTTTCAAGAGTGCCATGCCCGAAGGTTATGTACAGGAAAGAACTATATTTT<br>TCAAAGATGACGGGAACTACAAGACACGTGCTGAAGTCAAGTTTGAAGGTGATACCCTTGTTAATAGAATCGAG<br>TTAAAAGGTATTGATTTTAAAGAAGATGGAAACATTCTTGGACACAAATTGGAATACAACTATAACTCACACAA<br>TGTATACATCATGGCAGACAAACAAAAGAATGGAATCAAAGTTAACTTCAAAATTAGACACAACATTGAAGAT<br>GGAAGCGTTCAACTAGCAGACCATTATCAACAAAATACTCCAATTGGCGATGGCCCTGTCCTTTTACCAGACAA<br>CCATTACCTGTCCACACAATCTGCCCTTTCGAAAGATCCCAACGAAAAGAGAGACCACATGGTCCTTCTTGAGTT<br>TGTAACAGCTGCTGGGATTACACATGGCATGGATGAACTATACAAATAGGGATCCCATGGTACGCGT<br>CCTTCTCTCCT<br>CTGCAGCGGCCGCTACTAGTATTCATTCGACTATAACAAACCATTTTCTTGCGTAAACCTGTACGATCCTACAGG<br>T – 3’ |
| pTet+LacO – *gfp** – rrnBT1—*mCherry*— BioBrick Suffix - pLux | 5’ -<br>TCCCTATCAGTGATAGAGATTGACATCCCTATCAGTGATAGAGATACTGAGCACATCAGCAGGACGCACTGACC<br>GAATTCATTAAAGAGGAGAAAGGTACCTTGTGAGCGGATAACAAAAAGGCTTGTGAGCGGATAACAAACCTGG<br>GTCGACAAGGAGAGAAGGATGAGTAAAGGAGAAGAACTTTTCACTGGAGTTGTCCCAATTCTTGTTGAATTAGA<br>TGGTGATGTTAATGGGCACAAATTTTCTGTCAGTGGAGAGGGTGAAGGTGATGCAACATACGGAAAACTTACCC<br>TTAAATTTATTTGCACTACTGGAAAACTACCTGTTCCATGGCCAACACTTGTCACTACTTTCTCTTATGGTGTTCA<br>ATGCTTTTCAAGATACCCAGATCATATGAAACGGCATGACTTTTTCAAGAGTGCCATGCCCGAAGGTTATGTAC<br>AGGAAAGAACTATATTTTTCAAAGATGACGGGAACTACAAGACACGTGCTGAAGTCAAGTTTGAAGGTGATAC<br>CCTTGTTAATAGAATCGAGTTAAAAGGTATTGATTTTAAAGAAGATGGAAACATTCTTGGACACAAATTGGAAT<br>ACAACTATAACTCACACAATGTATACATCATGGCAGACAAACAAAAGAATGGAATCAAAGTTAACTTCAAAATT<br>AGACACAACATTGAAGATGGAAGCGTTCAACTAGCAGACCATTATCAACAAAATACTCCAATTGGCGATGGCCC<br>TGTCCTTTTACCAGACAACCATTACCTGTCCACACAATCTGCCCTTTCGAAAGATCCCAACGAAAAGAGAGACC<br>ACATGGTCCTTCTTGAGTTTGTAACAGCTGCTGGGATTACACATGGCATGGATGAACTATACAAACAGGGATCC<br>ATTTGTCCTACTCAGGAGAGCGTTCACCGACAAACAACAGATAAAACGAAAGGCCCAGTCTTTCGACTGAGCCT<br>TTCGTTTTATTTGGGATCCTTACTTGTACAGCTCGTCCATGCCGCCGGTGGAGTGGCGGCCCTCGGCGCGTTCGT<br>ACTGTTCCACGATGGTGTAGTCCTCGTTGTGGGAGGTGATGTCCAACTTGATGTTGACGTTGTAGGCGCCGGGCA<br>GCTGCACGGGCTTCTTGGCCTTGTAGGTGGTCTTGACCTCAGCGTCGTAGTGGCCGCCGTCCTTCAGCTTCAGCC<br>TCTGCTTGATCTCGCCCTTCAGGGCGCCGTCCTCGGGGTACATCCGCTCGGAGGAGGCCTCCCAGCCCATGGTCT<br>TCTTCTGCATTACGGGGCCGTCGGAGGGGAAGTTGGTGCCGCGCAGCTTCACCTTGTAGATGAACTCGCCGTCCT<br>GCAGGGAGGAGTCCTGGGTCACGGTCACCACGCCGCCGTCCTCGAAGTTCATCACGCGCTCCCACTTGAAGCCC<br>TCGGGGAAGGACAGCTTCAAGTAGTCGGGGATGTCGGCGGGGTGCTTCACGTAGGCCTTGGAGCCGTACATGA<br>ACTGAGGGGACAGGATGTCCCAGGCGAAGGGCAGGGGGCCACCCTTGGTCACCTTCAGCTTGGCGGTCTGGGT<br>GCCCTCGTAGGGGCGGCCCTCGCCCTCGCCCTCGATCTCGAACTCGTGGCCGTTCACGGAGCCCTCCATGTGCAC<br>CTTGAAGCGCATGAACTCCTTGATGATGGCCATGTTATCCTCCTCGCCCTTGCTCACCAT CCTTCTCTCCT<br>CTGCAGCGGCCGCTACTAGTATTCATTCGACTATAACAAACCATTTTCTTGCGTAAACCTGTACGATCCTACAGG<br>T – 3’ |

**Supplementary Table S3**: **mCherry sequences mutated to introduce antisense stop codons.** See Figure 4d. Note that all sequences are oriented 3’ to 5’ in order to show the introduction of stop codons at the mutation points (black-colored bases).

| | |
|---|---|
| *mCherry*-antisense stop codon 1<br>(mutation in black) | 3’-<br>TTACTTGTACAGCTCGTCCATGCCGCCGGTTGAGTGGCGGCCCTCGGCGCGTTCGTACTGTTCCACGAT<br>GGTGTAGTCCTCGTTGTGGGAGGTGATGTCCAACTTGATGTTGACGTTGTAGGCGCCGGGCAGCTGCA<br>CGGGCTTCTTGGCCTTGTAGGTGGTCTTGACCTCAGCGTCGTAGTGGCCGCCGTCCTTCAGCTTCAGCC<br>TCTGCTTGATCTCGCCCTTCAGGGCGCCGTCCTCGGGGTACATCCGCTCGGAGGAGGCCTCCCAGCCC<br>ATGGTCTTCTTCTGCATTACGGGGCCGTCGGAGGGGAAGTTGGTGCCGCGCAGCTTCACCTTGTAGAT<br>GAACTCGCCGTCCTGCAGGGAGGAGTCCTGGGTCACGGTCACCACGCCGCCGTCCTCGAAGTTCATCA<br>CGCGCTCCCACTTGAAGCCCTCGGGGAAGGACAGCTTCAAGTAGTCGGGGATGTCGGCGGGGTGCTTC<br>ACGTAGGCCTTGGAGCCGTACATGAACTGAGGGGACAGGATGTCCCAGGCGAAGGGCAGGGGGCCAC<br>CCTTGGTCACCTTCAGCTTGGCGGTCTGGGTGCCCTCGTAGGGGCGGCCCTCGCCCTCGCCCTCGATCT<br>CGAACTCGTGGCCGTTCACGGAGCCCTCCATGTGCACCTTGAAGCGCATGAACTCCTTGATGATGGCC<br>ATGTTATCCTCCTCGCCCTTGCTCACCAT- 5’ |
| *mCherry*-antisense stop codon 2<br>(mutation in black) | 3’-<br>TTACTTGTACAGCTCGTCCATGCCGCCGGTGGAGTGGCGGCCCTCGGCGCGTTCGTACTGTTCCACGAT<br>GGTGTAGTCCTCGTTGTGGGAGGTGATGTCCAACTTGATGTTGACGTTGTAGGCGCCGGGCAGCTGCA<br>CGGGCTTCTTGGCCTTGTAGGTGGTCTTGACCTCAGCGTCGTAGTGGCCGCCGTCCTTTAGCTTCAGCC<br>TCTGCTTGATCTCGCCCTTCAGGGCGCCGTCCTCGGGGTACATCCGCTCGGAGGAGGCCTCCCAGCCC<br>ATGGTCTTCTTCTGCATTACGGGGCCGTCGGAGGGGAAGTTGGTGCCGCGCAGCTTCACCTTGTAGAT<br>GAACTCGCCGTCCTGCAGGGAGGAGTCCTGGGTCACGGTCACCACGCCGCCGTCCTCGAAGTTCATCA<br>CGCGCTCCCACTTGAAGCCCTCGGGGAAGGACAGCTTCAAGTAGTCGGGGATGTCGGCGGGGTGCTTC<br>ACGTAGGCCTTGGAGCCGTACATGAACTGAGGGGACAGGATGTCCCAGGCGAAGGGCAGGGGGCCAC<br>CCTTGGTCACCTTCAGCTTGGCGGTCTGGGTGCCCTCGTAGGGGCGGCCCTCGCCCTCGCCCTCGATCT<br>CGAACTCGTGGCCGTTCACGGAGCCCTCCATGTGCACCTTGAAGCGCATGAACTCCTTGATGATGGCC<br>ATGTTATCCTCCTCGCCCTTGCTCACCAT- 5’ |
| *mCherry*-antisense stop codon 3<br>(mutation in black) | 3’-<br>TTACTTGTACAGCTCGTCCATGCCGCCGGTGGAGTGGCGGCCCTCGGCGCGTTCGTACTGTTCCACGAT<br>GGTGTAGTCCTCGTTGTGGGAGGTGATGTCCAACTTGATGTTGACGTTGTAGGCGCCGGGCAGCTGCA<br>CGGGCTTCTTGGCCTTGTAGGTGGTCTTGACCTCAGCGTCGTAGTGGCCGCCGTCCTTCAGCTTCAGCC<br>TCTGCTTGATCTCGCCCTTCAGGGCGCCGTCCTCGGGGTACATCCGCTCGGAGGAGGCCTCCCAGCCC<br>ATGGTCTTCTTCTGCATTACGGGGCCGTCGGAGGGGAAGTTGGTGCCGCGCAGCTTCACCTTGTAGAT<br>GAACTCGCCGTCCTGCAGGGATGAGTCCTGGGTCACGGTCACCACGCCGCCGTCCTCGAAGTTCATCA<br>CGCGCTCCCACTTGAAGCCCTCGGGGAAGGACAGCTTCAAGTAGTCGGGGATGTCGGCGGGGTGCTTC<br>ACGTAGGCCTTGGAGCCGTACATGAACTGAGGGGACAGGATGTCCCAGGCGAAGGGCAGGGGGCCAC<br>CCTTGGTCACCTTCAGCTTGGCGGTCTGGGTGCCCTCGTAGGGGCGGCCCTCGCCCTCGCCCTCGATCT<br>CGAACTCGTGGCCGTTCACGGAGCCCTCCATGTGCACCTTGAAGCGCATGAACTCCTTGATGATGGCC<br>ATGTTATCCTCCTCGCCCTTGCTCACCAT- 5’ |
| *mCherry*-antisense stop codon 4<br>(mutation in black) | 3’-<br>TTACTTGTACAGCTCGTCCATGCCGCCGGTGGAGTGGCGGCCCTCGGCGCGTTCGTACTGTTCCACGAT<br>GGTGTAGTCCTCGTTGTGGGAGGTGATGTCCAACTTGATGTTGACGTTGTAGGCGCCGGGCAGCTGCA<br>CGGGCTTCTTGGCCTTGTAGGTGGTCTTGACCTCAGCGTCGTAGTGGCCGCCGTCCTTCAGCTTCAGCC<br>TCTGCTTGATCTCGCCCTTCAGGGCGCCGTCCTCGGGGTACATCCGCTCGGAGGAGGCCTCCCAGCCC<br>ATGGTCTTCTTCTGCATTACGGGGCCGTCGGAGGGGAAGTTGGTGCCGCGCAGCTTCACCTTGTAGAT<br>GAACTCGCCGTCCTGCAGGGAGGAGTCCTGGGTCACGGTCACCACGCCGCCGTCCTCGAAGTTCATCA<br>CGCGCTCCCACTTGAAGCCCTCGGGGAAGGACAGCTTTAAGTAGTCGGGGATGTCGGCGGGGTGCTTC<br>ACGTAGGCCTTGGAGCCGTACATGAACTGAGGGGACAGGATGTCCCAGGCGAAGGGCAGGGGGCCAC<br>CCTTGGTCACCTTCAGCTTGGCGGTCTGGGTGCCCTCGTAGGGGCGGCCCTCGCCCTCGCCCTCGATCT<br>CGAACTCGTGGCCGTTCACGGAGCCCTCCATGTGCACCTTGAAGCGCATGAACTCCTTGATGATGGCC<br>ATGTTATCCTCCTCGCCCTTGCTCACCAT- 5’ |
| *mCherry*-antisense stop codon 5<br>(mutation in black) | 3’-<br>TTACTTGTACAGCTCGTCCATGCCGCCGGTGGAGTGGCGGCCCTCGGCGCGTTCGTACTGTTCCACGAT<br>GGTGTAGTCCTCGTTGTGGGAGGTGATGTCCAACTTGATGTTGACGTTGTAGGCGCCGGGCAGCTGCA<br>CGGGCTTCTTGGCCTTGTAGGTGGTCTTGACCTCAGCGTCGTAGTGGCCGCCGTCCTTCAGCTTCAGCC<br>TCTGCTTGATCTCGCCCTTCAGGGCGCCGTCCTCGGGGTACATCCGCTCGGAGGAGGCCTCCCAGCCC<br>ATGGTCTTCTTCTGCATTACGGGGCCGTCGGAGGGGAAGTTGGTGCCGCGCAGCTTCACCTTGTAGAT<br>GAACTCGCCGTCCTGCAGGGAGGAGTCCTGGGTCACGGTCACCACGCCGCCGTCCTCGAAGTTCATCA<br>CGCGCTCCCACTTGAAGCCCTCGGGGAAGGACAGCTTCAAGTAGTCGGGGATGTCGGCGGGGTGCTTC<br>ACGTAGGCCTTGGAGCCGTACATGAACTGAGGGGACAGGATGTCCCAGGCGAAGGGCAGGGGGCCAC<br>CCTTGGTCACCTTCAGCTTGGCGGTCTGGGTGCCCTCGTAGGGGCGGCCCTCGCCCTCGCCCTCGATCT<br>CGAACTCGTGGCCGTTCACTGAGCCCTCCATGTGCACCTTGAAGCGCATGAACTCCTTGATGATGGCC<br>ATGTTATCCTCCTCGCCCTTGCTCACCAT- 5’ |
| *mCherry*-antisense stop codon 6<br>BioBrick Suffix - pLux<br>(mutation in black, here in the BioBrick suffix) | 3’-<br>TTACTTGTACAGCTCGTCCATGCCGCCGGTGGAGTGGCGGCCCTCGGCGCGTTCGTACTGTTCCACGAT<br>GGTGTAGTCCTCGTTGTGGGAGGTGATGTCCAACTTGATGTTGACGTTGTAGGCGCCGGGCAGCTGCA<br>CGGGCTTCTTGGCCTTGTAGGTGGTCTTGACCTCAGCGTCGTAGTGGCCGCCGTCCTTCAGCTTCAGCC<br>TCTGCTTGATCTCGCCCTTCAGGGCGCCGTCCTCGGGGTACATCCGCTCGGAGGAGGCCTCCCAGCCC<br>ATGGTCTTCTTCTGCATTACGGGGCCGTCGGAGGGGAAGTTGGTGCCGCGCAGCTTCACCTTGTAGAT<br>GAACTCGCCGTCCTGCAGGGAGGAGTCCTGGGTCACGGTCACCACGCCGCCGTCCTCGAAGTTCATCA<br>CGCGCTCCCACTTGAAGCCCTCGGGGAAGGACAGCTTCAAGTAGTCGGGGATGTCGGCGGGGTGCTTC<br>ACGTAGGCCTTGGAGCCGTACATGAACTGAGGGGACAGGATGTCCCAGGCGAAGGGCAGGGGGCCAC<br>CCTTGGTCACCTTCAGCTTGGCGGTCTGGGTGCCCTCGTAGGGGCGGCCCTCGCCCTCGCCCTCGATCT |

| | |
|---|---|
| | CGAACTCGTGGCCGTTCACGGAGCCCTCCATGTGCACCTTGAAGCGCATGAACTCCTTGATGATGGCC ATGTTATCCTCCTCGCCCTTGCTCACCAT CCTTCTCTCCT CTGCAGCGGCCGCTACTACTATTCATTCGACTATAACAAACCATTTTCTTGCGTAAACCTGTACGATCC TACAGGT |

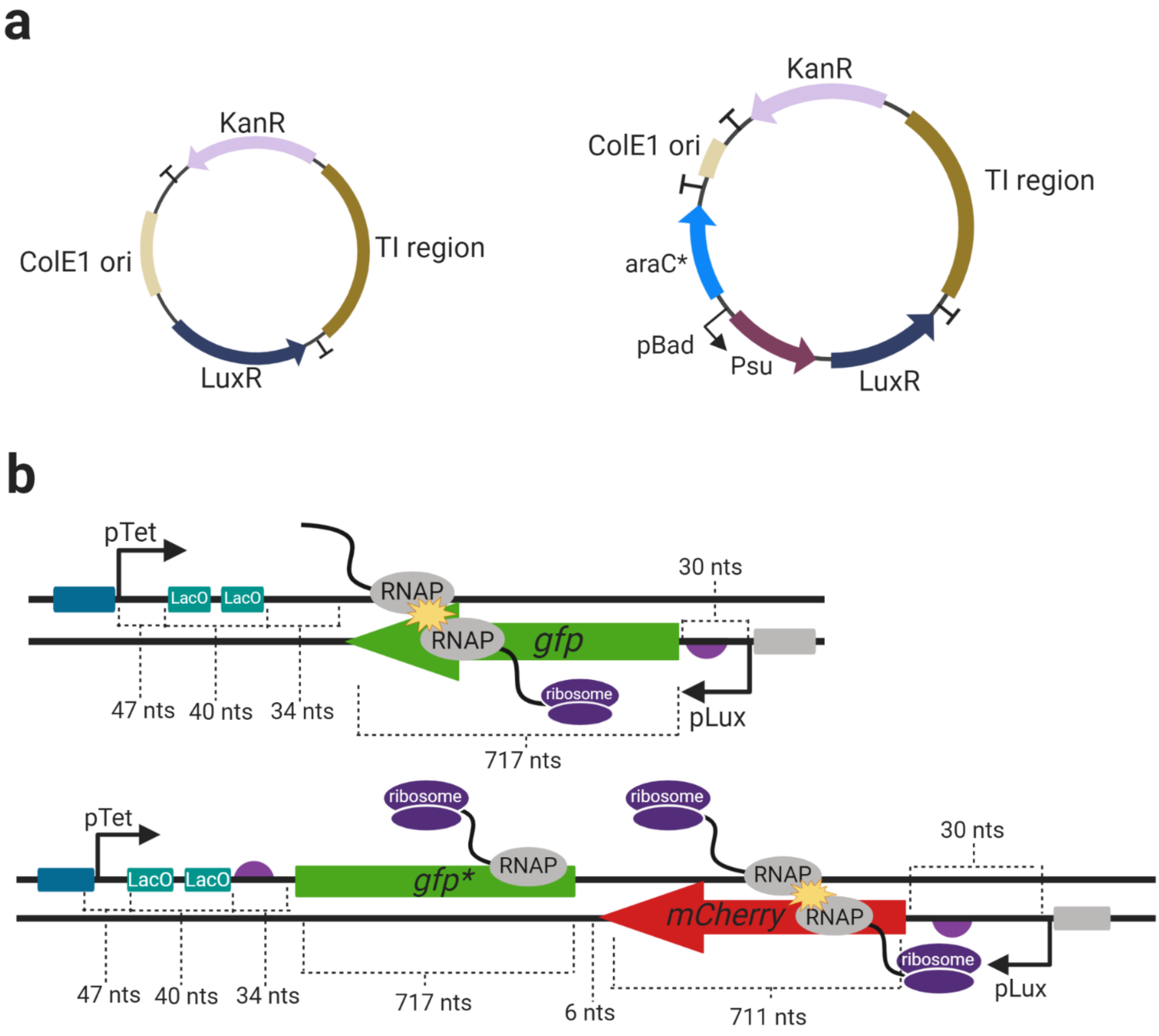


**Supplementary Figure S1**: **Maps of plasmids used in this study.** a) Plasmid maps showing the design of the constructs and do not contain (left, Figs. 1b-c, Fig. 2-4) and do contain (Figs. 1d-f and Supplementary Figure S3, S6) Psu and araC*. Note that features are not illustrated to scale. b) Diagrams of TI inserts showing the DNA lengths (in nucleotides) of different genetic features in the convergent promoter *gfp* (Fig. 1) and *gfp*-mCherry* (Figs. 2-4) constructs

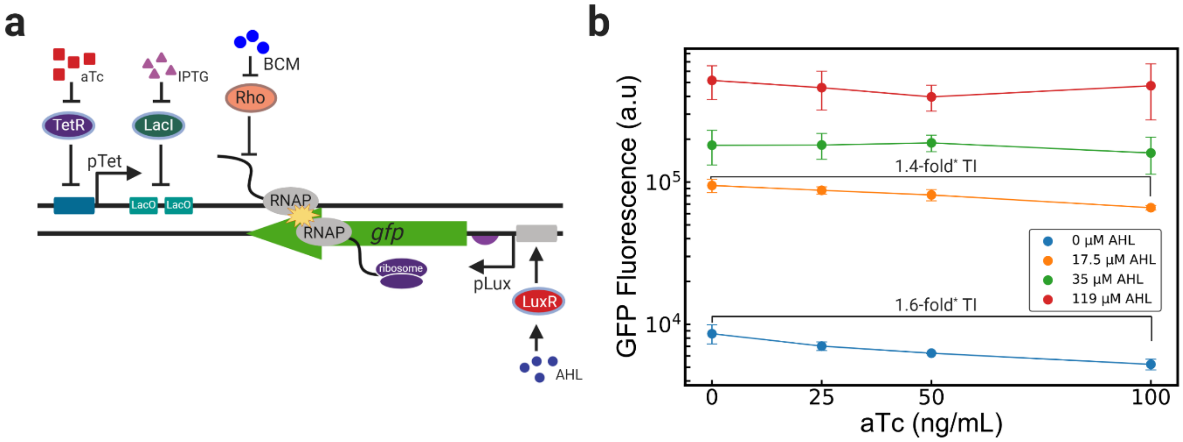


**Supplementary Figure S2**: **Convergent promoters without processivity control produces weak TI.** a) Diagram of inducible TI construct, also depicted in Fig 1b. b) Weak, aTc-dependent repression of GFP expression due to TI is observed at low AHL concentrations. All experiments were performed with 1 mM IPTG to remove the LacI roadblock. Error bars are denoted as ± s.d. Statistical significance was denoted as $p$<0.05, as determined through the Mann-Whitney $U$ test. n=3 biological replicates.

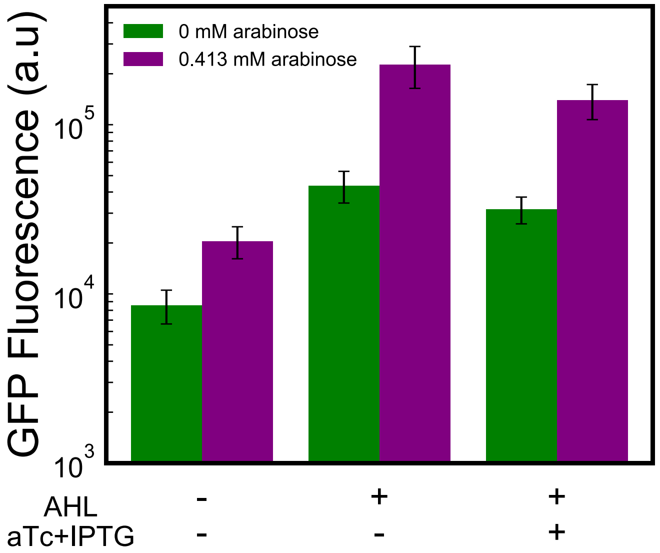


**Supplementary Figure S3: High Psu expression increases gene expression under all conditions.** GFP expression for the construct shown in Figure 2a with 0.413 mM arabinose in the presence of no inducers, 100 µM AHL only, and 100 µM AHL with 100 ng/mL aTc and 1 mM IPTG. Error bars are denoted as ± s.d. n=3 biological replicates.

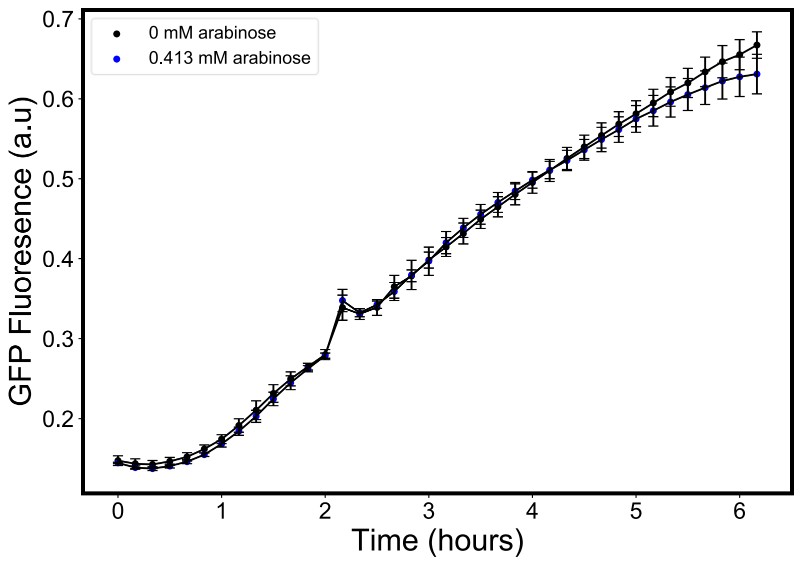


**Supplementary Figure S4: Psu expression exhibits negligible growth effects.** Growth curves for the construct shown in Figure 2a with and without 0.413 mM arabinose, in the absence of any AHL, aTc, or IPTG. Error bars are denoted as ± s.d. n=3 biological replicates.

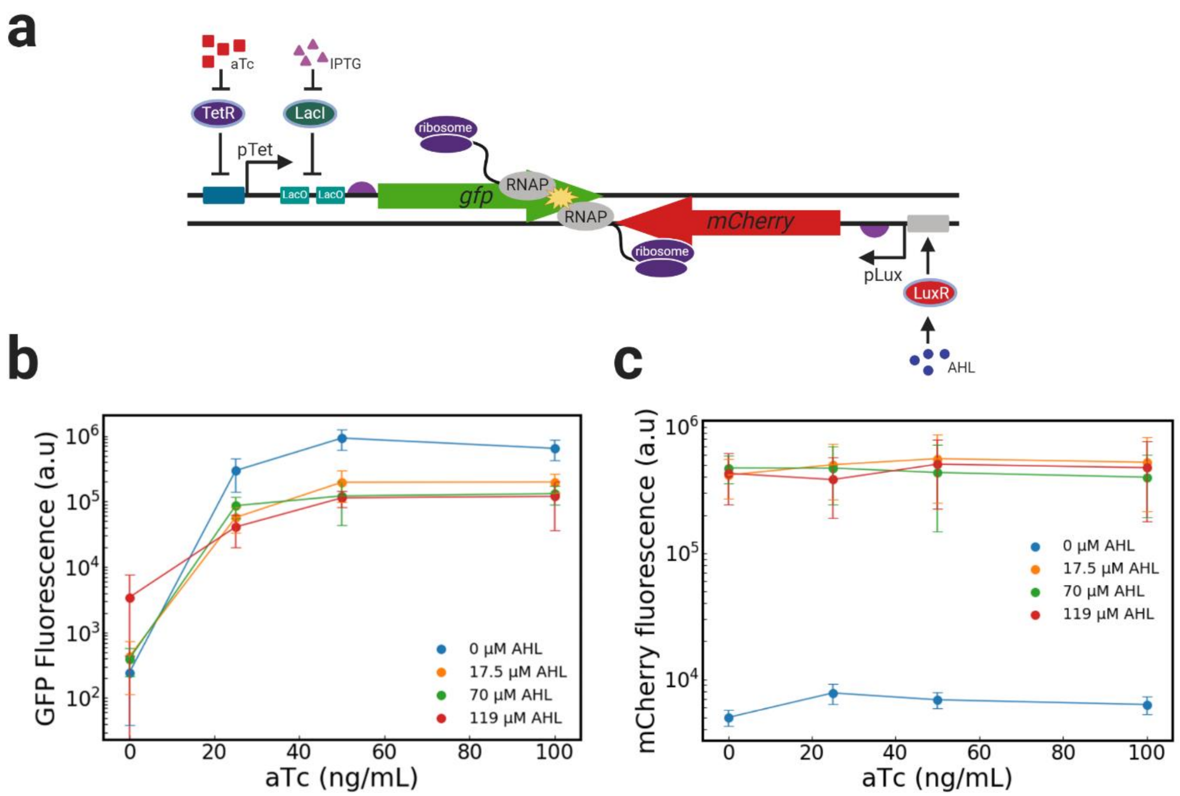


**Supplementary Figure S5: Reduction in GFP expression due to TI is observed in the *gfp-mCherry* convergent construct.** GFP fluorescence of the convergent *gfp-mCherry* construct as a function of both aTc and AHL. a) Diagram showing the *gfp-mCherry* construct. b) Approximately 7-fold TI for GFP expression is observed in the presence of saturating AHL concentrations. c) Negligible TI is observed for mCherry expression with increasing aTc. Error bars are denoted as ± s.d. Statistical significance was denoted as $p<0.05$, as determined through the Mann-Whitney $U$ test. n=3 biological replicates.

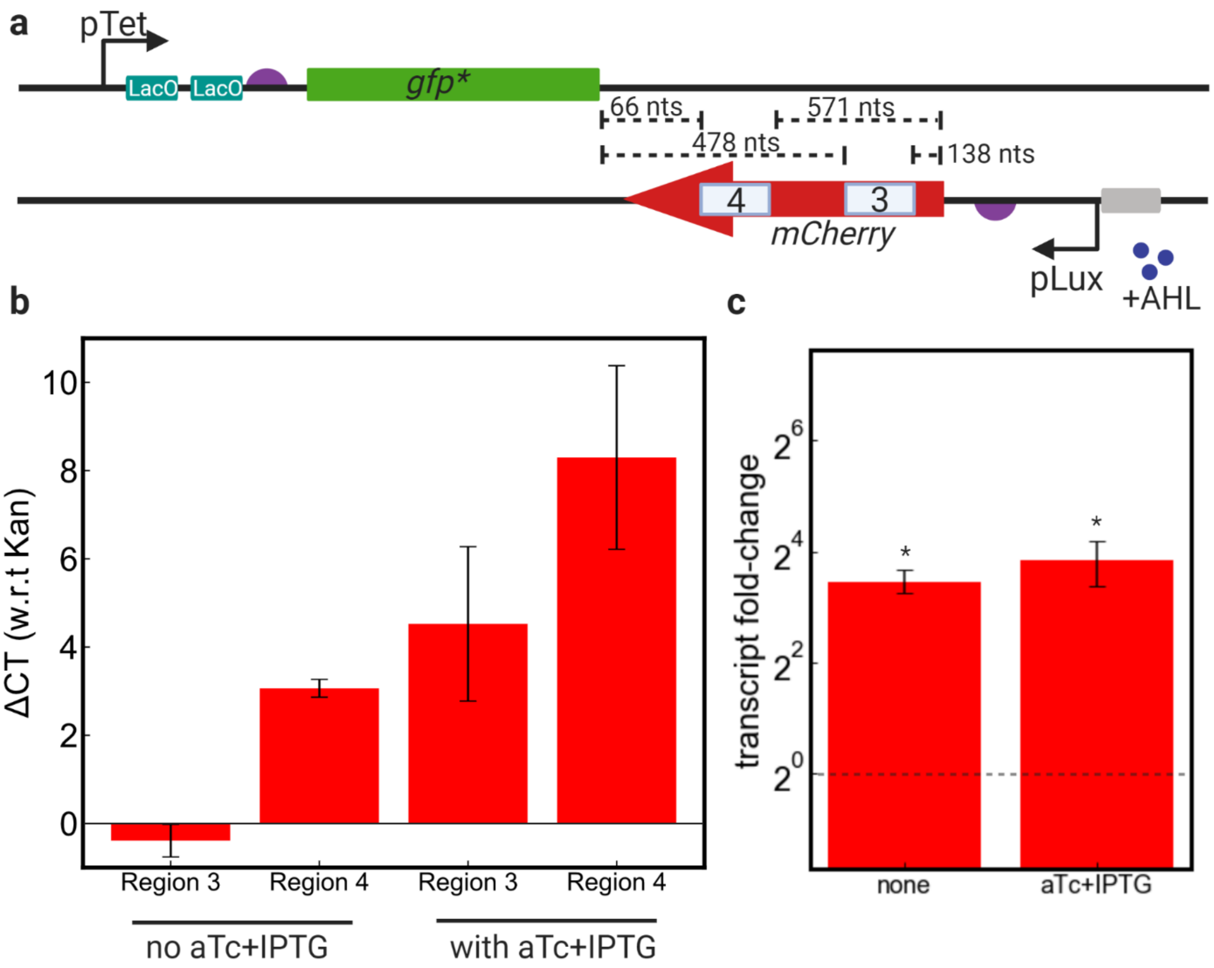


**Supplementary Figure S6: Truncated *mCherry* transcripts are present in the presence and absence of TI.** a) Strand-specific qPCR (Materials and Methods) was used to measure the abundance of *mCherry* transcripts containing either Region 3 or Regions 3 and 4 together in the *gfp*-mCherry* construct. b) $\Delta C_T$ with respect to *Kan* is reported for Region 3 and Region 4 in the presence and absence of interfering promoter activation (aTc+IPTG addition). c) The ~13-fold observed decrease in transcript abundance from Region 3 to Region 4 represents a ~13x more truncated transcripts (containing Region 3 and not Region 4) than transcripts containing both. Error bars are denoted as ± s.d, here represented as ($2^{-(\Delta\Delta CT+s.d)}$, $2^{-(\Delta\Delta CT-s.d)}$). * indicates significance (one-sample T-test, $p<0.05$) of the $\Delta\Delta C_T$ values with respect the null-hypothesis of $\Delta\Delta C_T=0$ (no change in transcript abundance between Region 3 and Region 4).

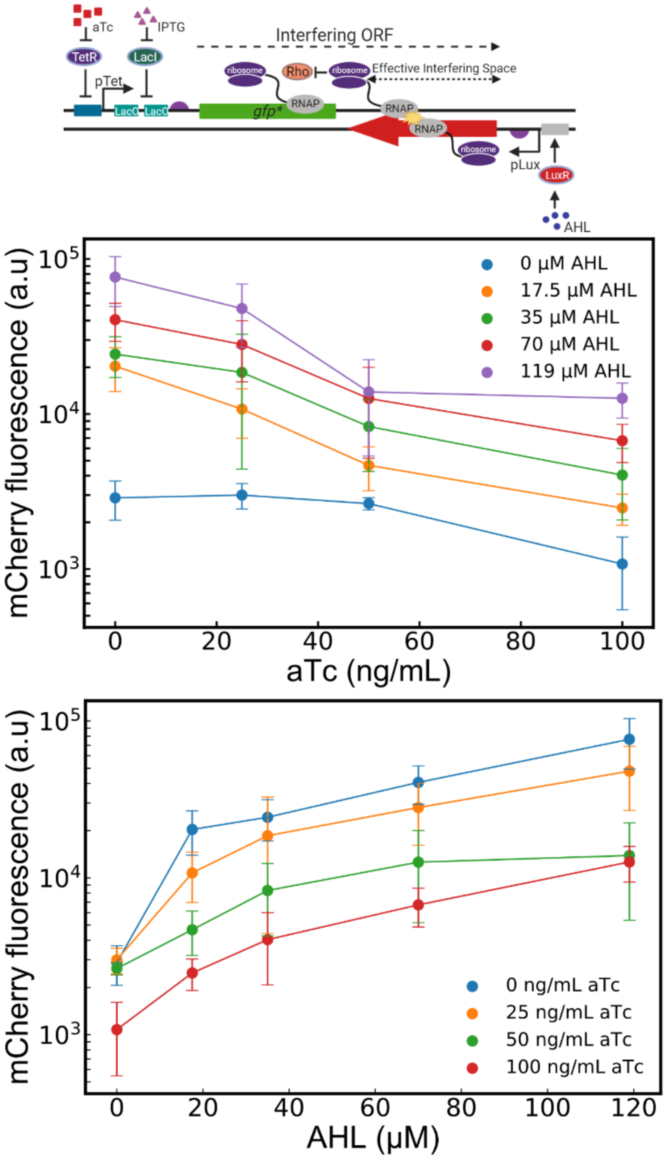


**Supplementary Figure S7: Expression control with processivity control yields a layered TI response.** Activation of the interfering promoter, pTet with different concentrations of aTc (at saturating 1mM IPTG to remove the LacI roadblock) and expressing promoter, pLux with different AHL concentrations tune the extent of TI. This interference and expression control coupled the ribosomal processivity control represent a layered response for TI-based genetic devices. For Fig. 2b, error bars are denoted as ± s.d. Statistical significance ($p$<0.05) was denoted with *, as determined through the Mann-Whitney $U$ test. n ≥ 3 for these experiments.

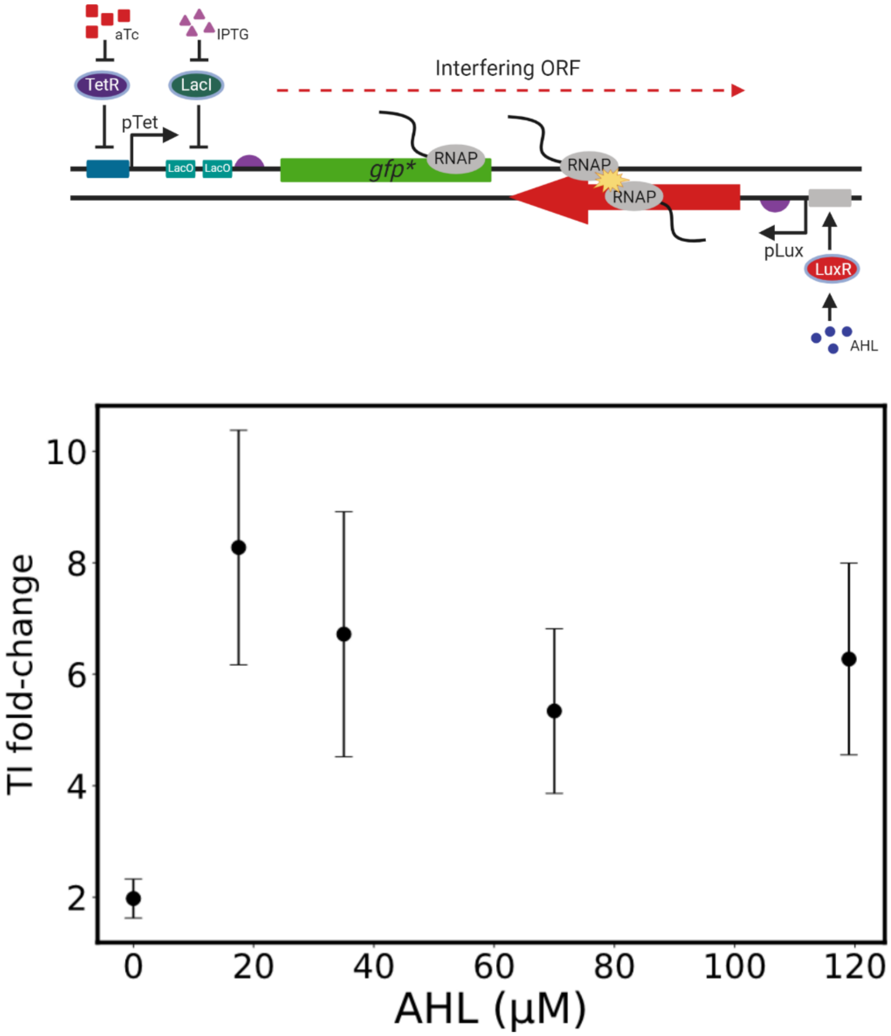


**Supplementary Figure S8: TI fold-change over the *gfp*-mCherry* is not a function of expressing promoter strength (AHL) when concentrations exceed 20 µL.** TI fold-change as a function of AHL concentration in the *gfp-mCherry* convergent construct (illustrated above). Fold-change is calculated by comparing AHL-only mCherry expression with mCherry expression in the presence of AHL with 100 ng/mL and 1 mM IPTG (Equation 1). Error bars are denoted as ± s.d. Statistical significance was denoted as $p<0.05$, as determined through the Mann-Whitney $U$ test. n=3 biological replicates.

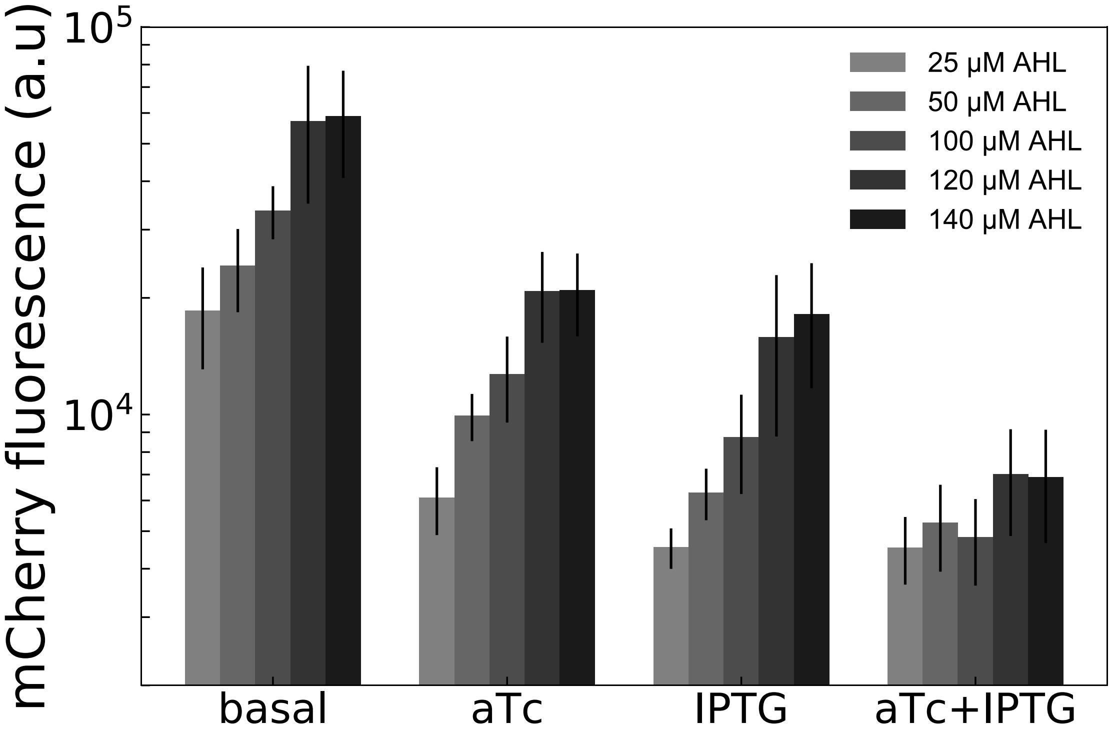


**Supplementary Figure S9: NOR behavior changes with expressing promoter strength.** Increasing AHL improves the fold-change of TI-repression (basal vs. aTc+IPTG) but also increases NOR asymmetry (aTc-only vs. IPTG-only vs. aTc+IPTG). Error bars are denoted as ± s.d. Statistical significance was denoted as $p<0.05$, as determined through the Mann-Whitney *U* test. n=3 biological replicates.
